# Divergent Electrophysiological Responses in the Human Hippocampus During Verbal Memory Processing

**DOI:** 10.64898/2026.01.28.702353

**Authors:** Weichen Huang, Julian Tobias Quabs, Dian Lyu, Eric K. van Staalduinen, Sofia Pantis, Olivia Marais, James Stieger, Eugene Liang, Gayle Deutsch, Zihuai He, Vivek Buch, Josef Parvizi

## Abstract

The profile of electrophysiological responses in the human hippocampus (HPC) during verbal memory processing has remained complex and unclear. Here, we studied 26 patients implanted with intracranial electrodes across 187 HPC sites (50% left, 2-18 per patient). During memory encoding and retrieval, a subset of HPC responsive sites demonstrated increased ripple events, along with elevated high-frequency (HFA >50 Hz), and low-frequency (LFA 1–8 Hz) activity. A nearly equal number of sites showed no changes in ripple rate but increased LFA power and a delayed response-locked decrease in HFA power. More importantly, both successful encoding as well as recognition of remembered words were strongly associated with the coordination of the timing of LFA and HFA increases across the two clusters of responsive HPC sites. Using direct cortical electrical stimulations, we confirmed overlapping, but partially distinct, cortical connections to the functionally distinct HPC clusters. Our findings suggest a mesoscale mosaic functional organization within the human HPC where adjacent sites with divergent electrophysiological responses may have specialized roles during verbal memory processing. More importantly, our findings suggest that successful human memory depends on the coordination of the timing of low and high frequency local fields generated across these functionally divergent neuronal population sites.

## INTRODUCTION

The hippocampus (HPC) is a critical hub for episodic memory, enabling event learning and memory that guides mnemonic decisions, such as whether a current stimulus was previously encountered or is novel (i.e., old-new recognition)^1^. Electrophysiological insights into human HPC functional organization have spanned spatial scales, including from the single unit level (micrometer resolution)^2^ to intracranial electroencephalography (iEEG)^3^ recordings of local field potentials (LFPs) at the mesoscopic level^4^ (millimeter resolution).

Extant studies have documented a distinct electrophysiological signature of activity within human HPC that is posited to play important roles in memory retrieval: Once the memory cue is presented, increased stimulus-locked theta (4–7Hz) and high-gamma activity (>50 Hz), increased ripple rates ^5–15^, along with reduced power in the mid-frequency (MFA,10-20Hz) range have been documented widely^16^. Moreover, interactions between low and high frequency activities have been documented in the form of classic coupling between the phase of theta and amplitude of high-gamma activity^17,18^.

Despite the great progress in direct electrophysiological recordings from the human HPC, the literature evidence has remained complex and at times confusing^12,19^. For instance, low frequency activity has encompassed not only slower (∼3Hz) but also faster (∼8Hz) activity across different subregions of the HPC^20^ - both of which fall *outside* the typical range of conventional theta band. Such findings have raised the possibility that “theta activity” in the human HPC during episodic memory processing–perhaps unlike the classic theta rhythm observed in rodents during spatial memory tests^21^ - may be a result of a spectral tilt and a part of a broader band of low frequencies^19^. Further complicating the picture, heightened high-gamma activity has been tied to an increase in the average firing rate of a population of neurons^22^, and thus it would be expected that successful memory processing (hallmarked by increased stimulus-locked HFA) must also be associated with increased firing rates of HPC neurons. Yet, direct neuronal recordings in the HPC have reported that only a fraction of HPC neurons increase their firing rate during memory retrieval, while a significant proportion of HPC neurons are indeed *deactivated* ^7,23–25^. In a recent study^26^, time-resolved representational similarity analysis of iEEG data from human HPC and various cortical regions revealed higher temporal similarity only at time points when the high gamma power was *reduced*. Together, these findings raise the possibility that decreases in hippocampal high frequency activity (HFA) may be a major part of episodic memory retrieval ^27^. Yet the issue of HFA deactivations during episodic memory processing has not been explored in detail.

Anatomical (e.g., ^28^) and neurophysiological (e.g., ^29^) studies in non-human (e.g., ^30^) and human (e.g.,^31^) brains, and recent findings from direct intracranial electroencephalography (iEEG) studies in the human brain^32–35^ have suggested a clear functional heterogeneity in the human brain at the mesoscopic level. For instance, a recent study uncovered that a distinct set of neuronal recording sites within the insula showed selective engagement during word encoding, while another set of sites (adjacent and interspersed with the memory-active sites) tracked stimulus valence. Electrical stimulation approach further demonstrated a selective, top-down causal influence from the memory- (but not the valence-) relevant insular sites to the ipsilateral HPC in the same subjects^35^. This finding highlighted an intricately heterogenous functional architecture at the mesoscopic scale, which is in line with a large body of intracranial EEG literature reporting selective and distinct profiles of electrophysiological responses across neighboring electrodes that are only millimeters apart^51^.

Given the extant evidence from human intracranial studies, together with emerging findings from non-human mammalian research demonstrating variations in anatomical connectivity and circuit architecture within the hippocampus (HPC), which may confer distinct functional roles to neighboring neuronal populations^36^, we hypothesized that a comparable mesoscopic-level functional heterogeneity exists in the human HPC. Such heterogeneity may not only account for, but also reconcile, the seemingly contradictory findings reported in the human hippocampal literature.

In the current study, we recruited 26 participants with temporal lobe epilepsy (TLE) who underwent iEEG recording during verbal encoding and old–new recognition task involving abstract and concrete words presented either minutes earlier (*immediate recognition*) or a day earlier (*delayed recognition*). We believe our findings help place prior literature on the hippocampus into a more coherent framework, particularly where earlier studies have yielded inconsistent or difficult-to-interpret results. Additionally, our study extends the recent findings of a “*salt-pepper functional organization*” at the millimeter scale in the human cerebral to the human HPC.

## RESULTS

### Participants

Our cohort consisted of 26 participants admitted for surgical evaluation of their refractory focal epilepsy (age range: 19-67 years old, 12 female). Only one participant was left-handed with bilateral language dominance. Demographic details are listed in Supplementary Table S1.

As part of routine clinical evaluation, participants were implanted with multiple electrodes for stereo-electroencephalography (sEEG) to localize the source of their seizures. We aggregated data from 187 sEEG recording sites across bilateral hippocampi (Figure 1a) (median: 6 sites per participant; range: 2-18 sites; left hemisphere: 94 sites, right hemisphere: 93 sites). Prior to participation in our experiments, all participants provided informed consent and volunteered their time for research. All procedures described in our study were approved by our university’s Institutional Review Board (IRB).

**Figure 1:**
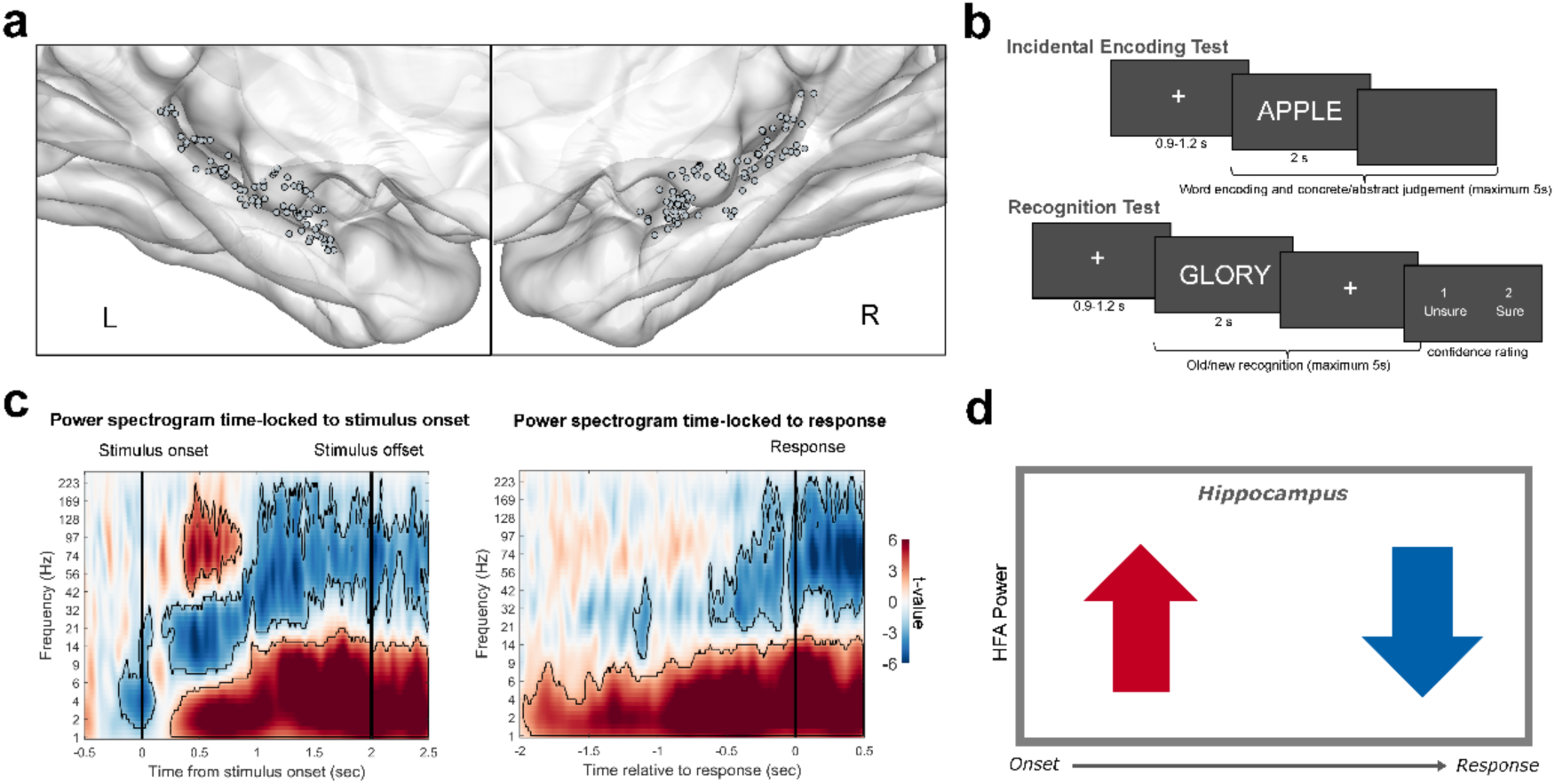
Electrophysiological Activity of Hippocampal Sites. **(a)** Electrode localization of 187 HPC sites across all 26 participants (left hemisphere: n = 94 sites, right hemisphere: 93 sites). **(b)** Experimental protocol of the memory task. The memory experiment included an incidental encoding test followed by an immediate recognition test on the first day and a delayed recognition test on the second day. During the incidental encoding test, participants were presented with 96 words, each displayed for 2 seconds. Participants were instructed to judge each word as either “concrete” or “abstract” and had up to 5 seconds to press a button to indicate their decision. In the recognition test, participants were shown another list of 96 words, consisting of 48 words from the incidental encoding phase (“old” words) and 48 new words. They were asked to classify each word as “new” or “old” within 5 seconds and to rate their memory confidence as low (“unsure”) or high (“sure”). **(c)** Power spectrograms of the hippocampus (HPC) during all good-performance blocks (161 sites) across immediate and delayed recognition tests, normalized relative to –500 to –200 ms baseline, and time-locked to stimulus onset (left) and response (right), revealed a sustained increase in low frequency activity (LFA) combined with an increase in high-frequency activity (HFA) following stimulus onset and a significant HFA decrease prior to the response. **(d)** The pattern in c could be interpreted as “*the signature of hippocampal response*” during recognition, as if the majority of HPC sites exhibit a stimulus-locked increase in HFA power followed by a response-locked decrease. Our further findings refuted this interpretation (see Figure 2).

### Behavioral Results

The memory experiment consisted of a word encoding task (96 trials), wherein participants classified individually presented words as abstract or concrete. This single encoding session was followed by a distracter task before an immediate and a 24-hr delayed old-new recognition test (96×2 = 192 trials); in both, they responded in a self-paced manner to individually presented word test probes, indicating if each was Old or New along with whether their decision confidence was high (sure) or low (unsure). (See Methods and Figure 1b).

We deliberately chose this well-established and relatively simple task to maximize comparability with the extant human memory literature. Our goal was not to introduce a novel behavioral paradigm, but rather to use a canonical task as a common framework to directly relate our neural findings to prior studies. By doing so, we aimed to bridge inconsistencies across the literature and reduce existing confusion regarding hippocampal function, thereby allowing new insight to emerge from neural heterogeneity rather than task complexity.

We gathered data regarding memory outcome (hit, miss, correct rejection, and false alarm trials), stimulus type (concrete vs. abstract words), experimental condition (immediate vs. delayed recognition), and judgment confidence (high vs. low). As expected, number of trials across memory outcomes varied depending on stimulus type, experimental condition, and judgment confidence (see details in Supplementary Information).

Our participants with chronic epilepsy, as expected^37^, had different degrees of memory performance (Supplementary Table S2) with different reaction times (RTs) (Supplementary Table S3). Specifically, regarding the memory performance, the D-prime values (0.71±0.57, median: 0.78), which reflects the ability to differentiate between new and old items (d′: z(hit rate) – z(false alarm rate)), revealed a better performance in immediate recognition (1.07±0.85, median: 1.19) than delayed recognition test (0.43±0.48, median: 0.40) (t(25) = 4.7337, p < 0.0001).

Reaction time (RT) during immediate recognition varied between memory conditions, wherein responses on hit trials (1.58±0.50s, median: 1.43s) were faster than on miss (1.90±0.71s, median: 1.74s; z = -3.5875, p = 0.0020), CR (1.87±0.58s, median: 1.71s; z = -3.7822, p = 0.0009) and false alarm trials (FA, 1.81±0.53s, median: 1.80s; z = -3.5875, p = 0.0020). Similar trends were observed during delayed recognition (hit: 1.95±0.76s, median: 1.70s; miss: 2.23±0.93s, median: 1.90s, z = - 1.8857, p = 0.0593; CR: 2.12±0.77s, median: 1.86s, z = -1.7838, p = 0.0745; FA: 2.07±0.83s, median: 1.87s, z = -1.5799, p = 0.1141). RTs for words recognized with high confidence (1.63±0.46s, median: 1.57s) were significantly faster than for those judged with low confidence (1.97±0.57s, median: 1.96s; z = -3.9494, p < 0.0001). Notably, the RTs did not differ between immediate recognition (1.74±0.53s, median: 1.61s) and delayed recognition (2.04±0.77s, median: 1.76s) (z = -0.5606, p = 0.5751) and did not differ between concrete (1.75±0.51s, median: 1.63s) and abstract trials (RT = 1.78±0.51s, median: 1.72s; z = -1.8668, p = 0.0619) (Supplementary Table S3 for details). In addition, we did not find any interactions on RTs between decision confidence and memory outcome (LMM, F = 0.6157, p = 0.6060), or between confidence and stimuli type (LMM, F = 0.0053, p = 0.9424).

As noted later, we mitigated the challenges posed by different number of trials in different memory and confidence conditions, and their interaction, by using i) trial-level linear mixed-effects model (LMM) analysis with subject and electrode as random effects, or ii) conducting analysis of data using only trials consisting of only one confidence type and one word stimulus type.

### Divergence of Electrophysiological Responses During Word Recognition

Our first analysis focused on the iEEG signature of recognition in the human HPC. For this, we analyzed data from immediate and delayed recognition trials in participants who performed with d-prime values higher than 0.5 (161 hippocampal recording sites, 22 participants with good performance in immediate recognition test and 10 participants with good performance in delayed recognition test). Coverage of electrodes in “good” performers is shown in Supplementary Figure S1.

During recognition trials when responses across all electrodes were pooled, the spectral EEG pattern showed the canonical signature that has been widely reported in prior studies, namely increased LFA (1–8 Hz), decreased MFA (10–40 Hz), and a marked rise in HFA (50–130 Hz, about 355–870ms after stimulus onset) followed by a reduction in HFA (about 630–80ms) before the subject’s response (Figure 1C). If one disregards mesoscopic-level functional heterogeneity within the HPC, the spectral EEG pattern shown in Figure 1c could be interpreted as *the signature of human HPC during memory retrieval* - as if all hippocampal sites contribute with the same electrophysiological responses by showing a stimulus-locked early increase of HFA power followed by a late response-locked decrease in the HFA power (Figure 1d). If this interpretation holds, HPC recording sites with early HFA increase should demonstrate a late decrease in their HFA power (i.e., the measure of stimulus-locked early HFA power and the late response-locked HFA power should be anti-correlated).

To test the above prediction, we first obtained trial-level measures of normalized HFA power for every recording site in a 600ms time window during the post-onset period (300ms to 900ms relative to stimulus onset) and a 600ms time window in the pre-response period (-650ms to -50ms relative to response), as both windows showed a significant HFA power change in the group-level analysis. Our analysis revealed an intriguing pattern (Figure 2a). Each site’s stimulus-locked HFA power did not anticorrelate with its own response-locked HFA power. Instead, the two HFA measures strongly correlated with each other (n = 161 electrodes, r = 0.6376, p < 0.0001). Only 3% (5/161) of HPC sites showed an early HFA increase followed by a late HFA decrease while 97% of HPC sites (156/161) did not. About 26% of HPC sites (N=42) showed no significant change of activity in either temporal windows. Of the responsive HPC sites, most (50 of 55, 91%) sites with statistically significant increase in stimulus-locked HFA activity showed *increased* (or no change in) response-locked HFA, and most sites (56 of 61, 92%) with significant response-locked HFA decrease had either decreased (or no change in) stimulus locked-HFA activity. Put simply, 74% of HPC sites showed significant changes of electrophysiological activity during recognition trials, and importantly, the profile of HFA changes across these sites was statistically neither random nor uniform (χ² = 52.6576, p < 0.0001). The location of the three groups of HPC sites in Figures 2a & 2b are shown as red, blue, and gray sites, respectively (Supplementary Table S4).

**Figure 2:**
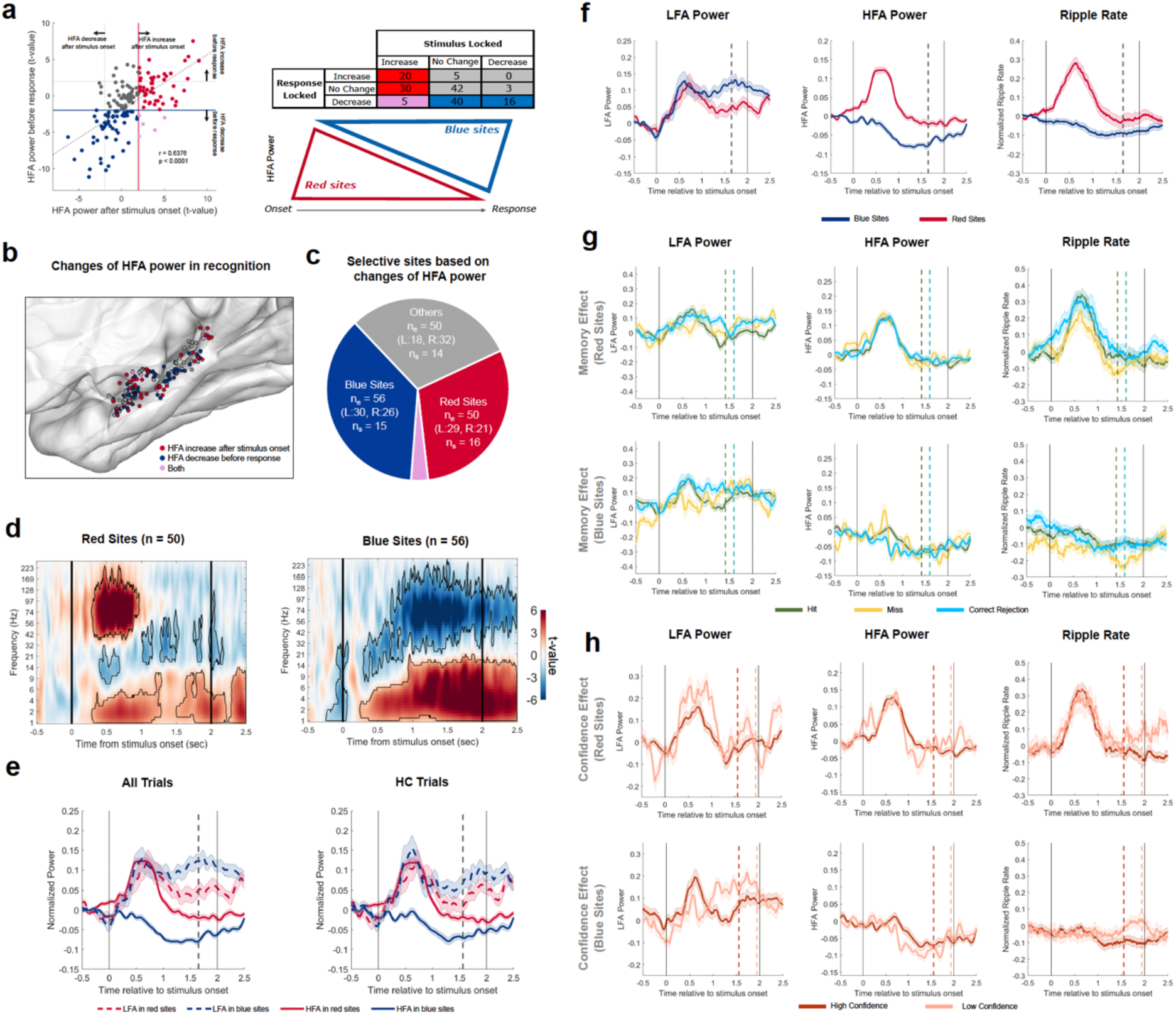
Distinct Functional Responses in Subsets of Hippocampal Sites. **(a)** HFA power in the post-stimulus and pre-response windows correlated directly as most of responsive sites with increased stimulus-locked HFA did not show decreased response-locked HFA (red sites), and most of responsive sites with decreased response-locked HFA did not show increased stimulus-locked HFA (blue). Only 3% of HPC sites exhibited both HFA increase after stimulus onset and HFA decreases before response (five purple sites). Each dot represents a distinct HPC site and the values for each site were calculated across all trials. The distribution of HFA changes across categories was not uniform (top right panel), indicating that the overall hippocampal profile as shown in Figure 1c was the summation of two subsets of sites with divergent gradient of responses, strong stimulus-locked increased HFA and strong response-locked decreased HFA (bottom right panel). **(b)** Sites showing HFA increases after stimulus onset (red sites) and those showing HFA decreases before response (blue sites) were distinct and exhibited a mosaic-like distribution throughout the HPC. All electrodes were projected to the right hemisphere. **(c)** Proportion of each site type among all HPC electrodes (ne: number of electrodes; ns: number of participants). **(d)** Power spectrograms time locked to stimulus onset for red sites (left) and blue sites (right). **(e)** Contrast of temporal dynamics in LFA and HFA in blue and red sites for all experimental trials (left panels) and all high-confidence (HC) trials (right panels). **(f)** Temporal dynamics of normalized low-frequency activity (LFA) power (left), HFA power (middle), and ripple rate (right) time-locked to stimulus onset for red and blue sites. **(g)** Comparisons of normalized LFA power (left), HFA power (middle), and ripple rate (right) timeseries time-locked to stimulus onset for red sites (top) and blue (bottom) sites across different memory outcomes for high-confidence (abstract and concrete) trials. **(h)** Comparisons of normalized LFA power (left), HFA power (middle), and ripple rate (right) timeseries time-locked to stimulus onset for red sites (top) and blue sites (bottom) across different confidence judgement for hit (concrete and abstract) trials. The dashed vertical lines in e, f, g, h represent the median reaction time for specific condition across all participants.

Further analyses confirmed that the HFA power in the red sites was comparable to the HFA power in the blue sites at pre-stimulus baseline (t(104) = −1.84, p = 0.0688). As shown in Figure 2f, HFA power at blue sites remained unchanged relative to baseline for several hundreds of milliseconds before subsequently exhibiting a significant decrease suggesting that the profile of stimulus-evoked responses in these sites cannot be explained by a mere stimulus-evoked reduction of activity from baseline. Similarly, the stimulus evoked activity at red sites initially increased during the recognition phase but started decreasing around the same time as the HFA power decreased at blue sites and returned to a level comparable to baseline. Moreover, the opposite changes of activity were not the same across memory conditions. Thus, the observed changes of HFA power during recognition trials reflected differential electrophysiological responses during the trial rather than differences in the pre-trial baseline activity.

Compared to red and blue sites, the gray sites demonstrated significantly higher pathological epileptic activity in the form of pathological high frequency oscillations (pHFOs) (LMM: F = 6.2560, p = 0.0135; Wilcoxon rank-sum test: z = 2.6081, p = 0.0091) and showed a similar trend in interictal epileptiform spikes (LMM: F = 3.1093, p = 0.0799; Wilcoxon rank-sum test: z = 2.1163, p = 0.0343). Furthermore, red sites (χ² = 1.6324, p = 0.2014) and blue sites (χ² = 0.3115, p = 0.5768) were found in both hippocampi without any lateralization effect, whereas grey sites (i.e., those without significant changes) were predominantly located in the right hemisphere (χ² = 5.5571, p = 0.0184).

Importantly, as detailed in Supplementary Tables S5 and S6, our findings revelaed that red and blue sites are intermingled throughout the HPC - rather than anatomically segregated or topographically ordered. Blue and red sites did not show any localization effects in any specific HPC subregions or subfields in the left hemisphere - though, in the right hemisphere, more red sites were located in the hippocampal tail (χ² = 8.5157, p = 0.0035) and CA1 subregions (χ² = 8.5800, p = 0.0034), and more blue sites were located in the hippocampal body (χ² = 9.9472, p = 0.0016) and CA4/DG subfields (χ² = 7.4529, p = 0.0063). We did not have sufficient sites in CA2/CA3 subfields (only 4 electrodes).

In the analysis of ripples, we were cognizant of the challenges in differentiating ripple activity from other high frequency activities^38^, and thus used a stringent method to identify ripples and differentiating them from broad HFA activity (Supplementary Figure S2). However, we are mindful that the measure of HFA power could clearly be affected by the presence or absence of ripples. Bearing this limitation in mind, we found that the ripple rates, during retrieval condition, increased only in the red sites (F = 19.5190, p = 0.0002) but *decreased* in the blue sites (F = 10.4556, p = 0.0062). The grey sites did not exhibit significant changes in either HFA power (F = 0.7280, p = 1) or ripple rate (F = 0.7807, p = 1). (Supplementary Figure S3). Notably, the ripple rate in baseline did not differ between different groups of sites (red sites: 0.34 ± 0.15 events/sec, blue sites: 0.39±0.12 events/sec, grey sites: 0.36±0.18 events/sec, F = 1.5710, p = 0.2112), demonstrating all hippocampal sites generated ripples, but showed different changes during the recognition trials.

In both red and blue site groups, we recorded an overall increase in LFA power (Supplementary Figure S3; LMM: red, F = 12.5327, p = 0.0027; blue, F = 9.2174, p = 0.0037; all p-values Bonferroni corrected across the three groups), but grey sites did not show this change (LMM, F = 4.8269, p = 0.0983). However, the red sites showed a significantly stronger LFA increase during the post-onset period (F = 11.9131, p = 0.0035), while both blue and red sites showed LFA increases during the pre-response period (LMM; red, F = 10.7378, p = 0.0058; blue, F = 16.7894, p = 0.0004).

Given that changes in the LFA and HFA are rarely isolated to a narrow high frequency band of EEG signal, but instead represent a spectral tilt or change in the exponent of the 1/f power of the recorded electrophysiological signal^39–43^, we measured the exponent of the 1/f signal (Supplementary Figure S4) as well as the event-related potentials (ERP) across the subtypes of the HPC sites. At group level, the blue sites exhibited a significant increase of aperiodic exponent compared to the baseline (LMM, F = 13.6134, Bonferroni-corrected p = 0.0015), while the red sites did not show any significant changes in the exponent (F = 2.1545, p = 0.4456). Additionally, the proportion of sites with an increase in the exponent of aperiodic activity was significantly higher in blue sites than in red sites (χ² = 8.3286, p = 0.0039), while the proportion of sites with an exponent decrease was significantly higher in red sites (χ² = 4.6557, p = 0.0310) suggesting that the changes in the HFA activity may also be accompanied by changes in a broad range of lower frequency bands of activity as well.

As for the event related potentials (ERPs), they did not reach significance at group level compared to the baseline (0-500ms relative to the stimulus onset) in either red sites (LMM, F = 4.7932, Bonferroni-corrected p = 0.1001) or blue sites (LMM, F = 4.2521, p = 0.1318). Moreover, we did not find consistent changes of ERP with changes in either LFA or HFA power.

### Memory Outcome and Changes in LFA and HFA Activity and Ripple Rates

The observation of varying gradients in HPC responses during recognition trials (i.e., heterogeneity of electrophysiological responses within the HPC) prompted us to investigate the possibility of *functional* heterogeneity within the HPC at the mesoscopic (millimeter) scale. Put simply, we predicted to find strong association between changes in the power of activity and memory outcomes.

We focused on the two groups of sites showing the most divergent profiles (red and blue sites) in terms of the power of HFA during recognition trials. Throughout the text, for the sake of simplicity, we refer to these sites as simply “red and blue sites” or “sites with divergent responses”.

We examined changes in their LFA and HFA power, along with ripple rates, across different memory conditions. We analyzed how the profile of these three classic neural markers of memory processing varied as a function of experimental factors: immediate vs. delayed recognition, word type (concrete vs. abstract), memory outcome (hits vs. correct rejections (CR)), and judgement confidence (high vs. low) (Figure 2f-h and Supplementary Figure S5). We had an insufficient number of miss and false alarm trials for statistical comparisons; nevertheless, we illustrate the data from miss trials in figures visual comparison may be helpful.

To mitigate the challenges posed by different number of trials in different memory and confidence conditions, and given the interaction between them, we employed two complementary approaches. First, we conducted a trial-level linear mixed-effects model (LMM) analysis to examine potential confounding effects and interactions of memory outcome, stimulus type, and judgment confidence within red and blue sites, focusing on LFA power, HFA power, and ripple rates during the post-onset period. As this approach revealed significant effects and confounding effects for judgment confidence (high vs. low confidence) and stimulus types (concrete vs. abstract words), we selected a second approach by focusing on high confidence concrete trials (Supplementary Figure S5 and Supplementary Table S7).

To control for effects of memory confidence, we examined the effects of memory outcome and stimulus type by focusing exclusively on high-confidence trials, which were significantly more frequent than low-confidence trials (65.42% ± 21.28%, Wilcoxon signed-rank test, z = 2.9057, p = 0.0037). We found significant increases in LFA power (F = 10.1708, p = 0.0025), HFA power (F = 61.8952, p < 0.0001), and ripple rate (F = 23.0922, p < 0.0001) in red sites in not only hits but also correct rejection (CR) trials (Supplementary Table S8). Changes of activity in red sites did not differ across different memory conditions (LFA: F = 2.4398, p = 0.1184; HFA: F = 0.7121, p = 0.3988; Ripple: F = 0.7467, p = 0.3876), suggesting that the strong changes of activity in red HPC sites was generically present when the subject was involved in retrieving memories, but the power of these changes did not predict successful retrieval. By contrast, memory effects could be observed in blue sites, with significant effects associated with changes in LFA (F = 8.3813, p = 0.0038) as well as HFA (F = 10.1110, p = 0.0015) power in these sites. In terms of ripples, there were no significant differences in their rate between hit and CR trials in either blue (F = 0.0676, p = 0.7949) or red sites (F = 1.0316, p = 0.3099).

LFA and HFA power in both blue and red sites was significantly higher for concrete than abstract stimuli [LFA: (LMM; blue: F = 32.7211, p < 0.0001; red: F = 31.3269, p < 0.0001); HFA: (LMM; blue: F = 6.5580, p = 0.0105; red: F = 6.9482, p = 0.0084)]. Additionally, both red and blue sites showed higher LFA power for immediate than delayed recognition trials (LMM; blue: F = 18.0586, p<0.0001; red: F = 8.5462, p = 0.0035); however, red sites exhibited higher HFA power during delayed recognition trials (F = 26.8633, p <0.0001). As shown in Supplementary **Figure S5**, we also analyzed memory effects and the temporal dynamics of changes in HFA and LFA during either only immediate or only delayed recognition trials, or either only concrete or only abstract trials to eliminate the confounding effects of interactions. These analyses confirmed again that blue sites had distinct changes in activity across different memory conditions, whereas the significant changes in LFA, HFA power, and ripple rates in red sites did not differentiate across memory outcomes. These conclusions were further confirmed by additional analyses, that was conducted including all 4 memory outcomes (hit, miss, CR and FA) using data from both immediate and delayed recognition of all 26 patients (see Supplementary Information).

### Memory Outcomes and Coordination of Activity Across HPC Sites

The above findings pointed to distinct electrophysiological profiles across red and blue HPC sites, possibly reflecting different functional roles in memory retrieval. Yet, although changes in the blue sites showed statistically significant associations with memory outcomes (Supplementary Table S7), they indicated that rises in LFA and reductions in HFA power only in blue sites were more pronounced during hits trials, and yet the HFA power and ripple rates in red or blue sites were no different during the hit trials. As seen in Figure 2g, we were surprised by the subtlety of the effects in blue sites and equally puzzling was the absence of any link between memory outcomes and the very pronounced increases in HFA and ripple activity observed at red sites.

Given that the red and blue sites were found adjacent to each other and distributed across the hippocampal subfields and subregions, we hypothesized that the *coordinated timing of activity* across these discrete HPC sites, instead of the *power of activity in discrete bands*, may serve as a stronger electrophysiological correlate of successful memory retrieval. Put simply, instead of changes in the power of activity in a given group of HPC sites, the synchronization of activity across distinct sites may be a more meaningful indicator of how information is integrated across distinct population of neurons within the HPC during retrieval.

To this end, we first analyzed the time of onset of increased HFA activity in red sites compared to the time of onset of HFA decreases in the blue sites using data recorded simultaneously from red and blue sites in the same hemisphere and in the same individual brains. This analysis revealed a significant temporal relationship between the activity of red and blue sites: increases in the HFA power at red sites occurred significantly before the decreases in HFA power at blue sites (red sites: 0.64s ± 0.18s vs. blue sites: 0.89s ± 0.22; LMM, F = 53.44, p < 0.0001).

Next, we analyzed the timing of changes in LFA and HFA across blue and red electrodes. We found significant synchronization of HFA in red sites with LFA in either red or blue sites that differed as a function of memory performance: such synchronization was present only during hit trials and was absent in miss or correct-rejection trials (Figure 3). The correlation coefficient index within the entire trial window between HFA in red sites and LFA in blue sites showed a significant effect of memory outcome (F = 12.098, p < 0.0001). Specifically, the index for hit trials was significantly higher than that for miss (t = 4.826, p < 0.0001) and CR (t = 3.235, p = 0.0014) trials. Similarly, the index between LFA in red sites and HFA in red sites also varied across memory outcomes (F = 3.7410 p = 0.0243), with hit trials showing significantly higher indices than miss (t = 2.735, p = 0.0192) trials (Supplementary Figure S6a). By contrast, the correlation index between LFA in red sites and LFA in blue sites did not exhibit memory outcome (F = 1.1131, p = 0.3305).

**Figure 3:**
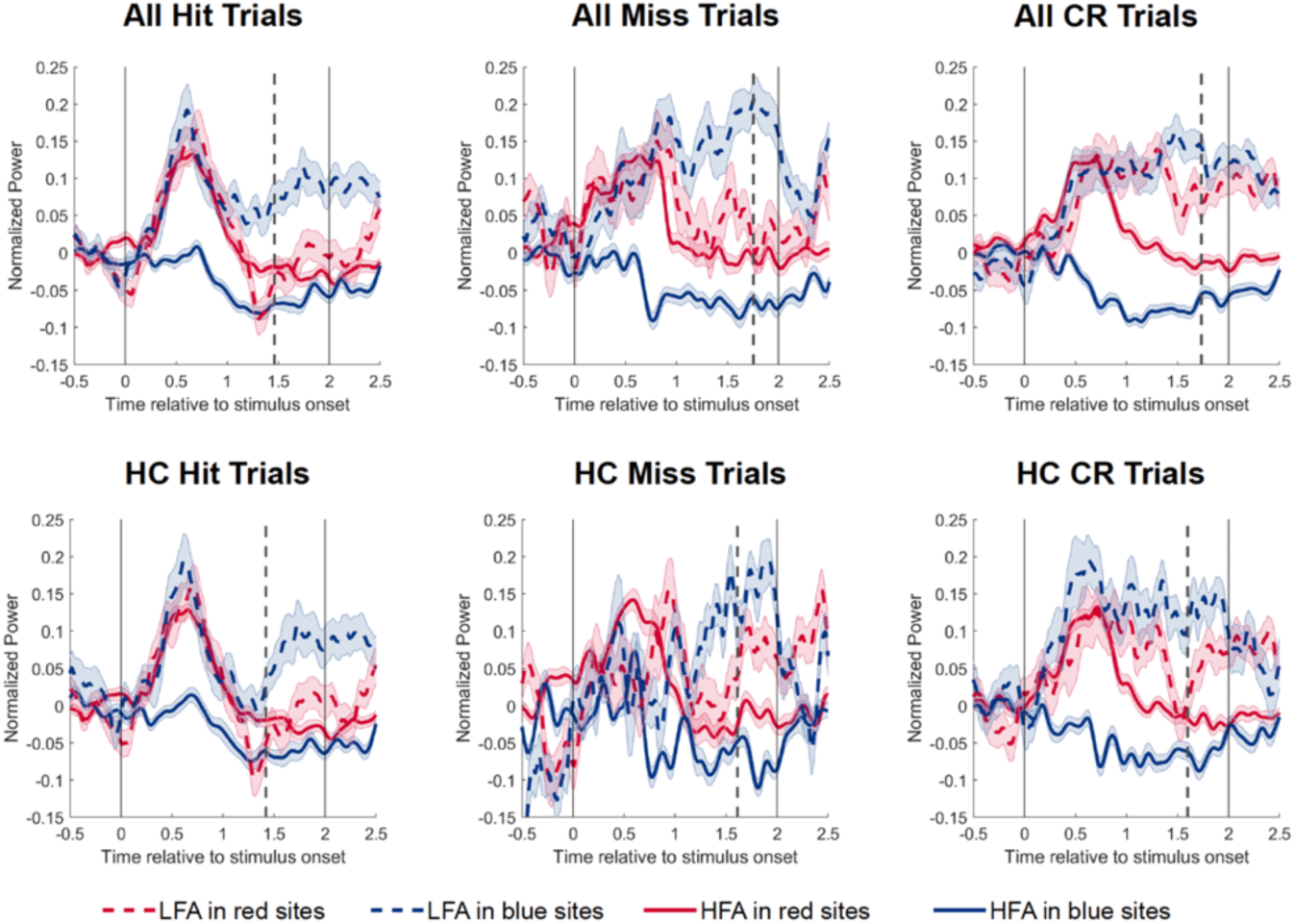
Memory Outcomes and Coordination of Activity Across HPC Sites. Hit trials are hallmarked by synchronized rise and fall of HFA (solid lines) and LFA (dashed lines) across sites with divergent HFA responses (red and blue lines referring to red and blue sites, respectively). High confidence trials are hallmarked by the fall of LFA (specially in blue sites) synchronized with the fall of other low and frequency activities. CR: correct rejection; HC: High confidence. Only data from electrodes with a minimum of five trials per condition were included in the group average.

Above findings were replicated by controlling for the effect of RT. We used a fixed time window (from stimulus onset to the median RT) and re-calculated the connectivity index. Once again, the index differed across memory outcomes (LMM; red LFA–red HFA: F = 3.2133, p = 0.0409; blue LFA–red HFA: F = 14.501, p < 0.0001). These results indicated that a coordinated activity across low and high frequency activities in blue and red sites underlies successful retrieval.

Lastly, we confirmed significant phase–amplitude coupling (PAC) ^18,44^ between theta phase and HFA amplitude (300–900 ms post-stimulus) in both blue and red sites similarly (LMM, F = 0.50, p = 0.48). PAC across ipsilateral site pairs was the same if the theta phase was chosen in blue or red sites (F = 0.06, p = 0.87).

### HPC Profiles During Encoding

We conducted post hoc analyses to examine the responses of different HPC neuronal populations during encoding. When grand-averaged across all HPC sites, we observed a significant increase in LFA and HFA, accompanied by a decrease in MFA, following stimulus onset (cluster-based permutation test, p < 0.05). However, red and blue sites continued to exhibit distinct response profiles (Figure 4a). Specifically, red sites showed a significant increase in HFA power (LMM, F = 4.7518, p < 0.0001) and ripple rate (LMM, F = 5.2723, p < 0.0001), whereas blue sites did not show significant changes in either HFA power (LMM, F = 0.7728, p = 0.4429) or ripple rate (LMM, F = 1.9114, p = 0.0611) after stimulus onset. In contrast, both red (F = 4.0036, p = 0.0002) and blue sites (F = 2.6276, p = 0.0111) exhibited significant increases in LFA (Figure 4b). With respect to memory effects, neither HFA power, LFA power, nor ripple rate differed between encoded and forgotten trials in either red or blue sites (Figure 4c). We also measure the correlation of LFA and HFA dynamic in blue and red sites during encoding. Similar with the recognition, the correlation coefficient index within the entire trial window between HFA in red sites and LFA in both blue (LMM, F = 11.881, p = 0.0007) and red (LMM, F = 9.5953, p = 0.0021) sites showed a significant effect of memory outcome, with encoded trials showing significantly higher indices than forgotten trials (Supplementary Figure S6b). By contrast, the correlation index between LFA in red sites and LFA in blue sites did not exhibit memory outcome (LMM, F = 0.0992, p = 0.7532).

**Figure 4:**
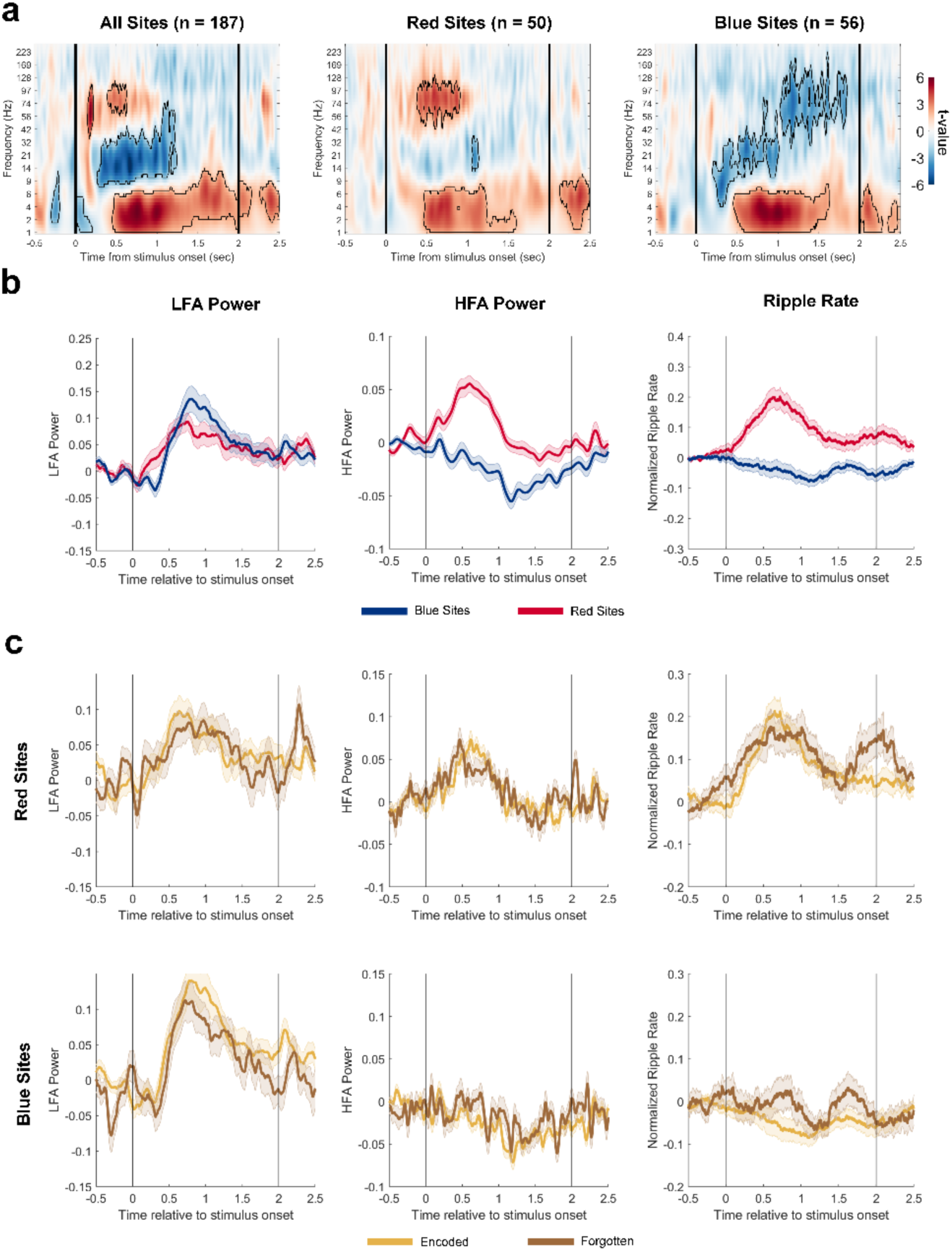
HPC Profiles During Encoding Task. (**a**). Power spectrum of all HPC sites, as well as red and blue sites during memory encoding. In the spectrograms, time–frequency clusters showing significant power increases (red) or decreases (blue) are outlined (p < 0.05, cluster-based permutation tests). (**b**) Temporal dynamics of normalized low-frequency activity (LFA) power (left), HFA power (middle), and ripple rate (right) time-locked to stimulus onset for red and blue. (**c**) Comparisons of normalized LFA power (left), HFA power (middle), and ripple rate (right) timeseries time-locked to stimulus onset for red sites (top) and blue (bottom) sites between encoded and forgotten trials during encoding task. Vertical black lines indicate stimulus onset and offset.

### Causal Connections of Red and Blue Sites with Distinct Anatomical Regions

As the process of retrieval process depends critically on the interaction between cortical regions and the HPC^45^, we examined if cortical inputs to the red and blue HPC sites were identical or distinct. We used data from direct single-pulse electrical stimulations across widespread brain regions while recording evoked responses in each HPC recording site —a method widely known as the cortico-cortical^46^ evoked potentials (CCEP) approach. We identified 2,519 stimulation–recording pairs involving red sites (11 participants) and 2,993 such pairs involving blue sites (9 participants). To quantify the strength of direct connections from each stimulation site to the HPC recording site, we computed the F1 index, as described in a recent study^47^. We divided the electrodes into different parcellations according to the Julich Brain Atlas^48^ and analyzed the connectivity flow from each cortical parcellation to the HPC sites.

In our analysis, we first quantified the strength of causal connections between the cortical parcels where we had implanted electrodes and the identified blue or red HPC sites. Stimulation of many cortical parcels evoked electrophysiological changes in the red or blue sites. However the strength of causal connections seemed to be different across cortical regions (darker red or blue colors in Figure 5a & b).

**Figure 5:**
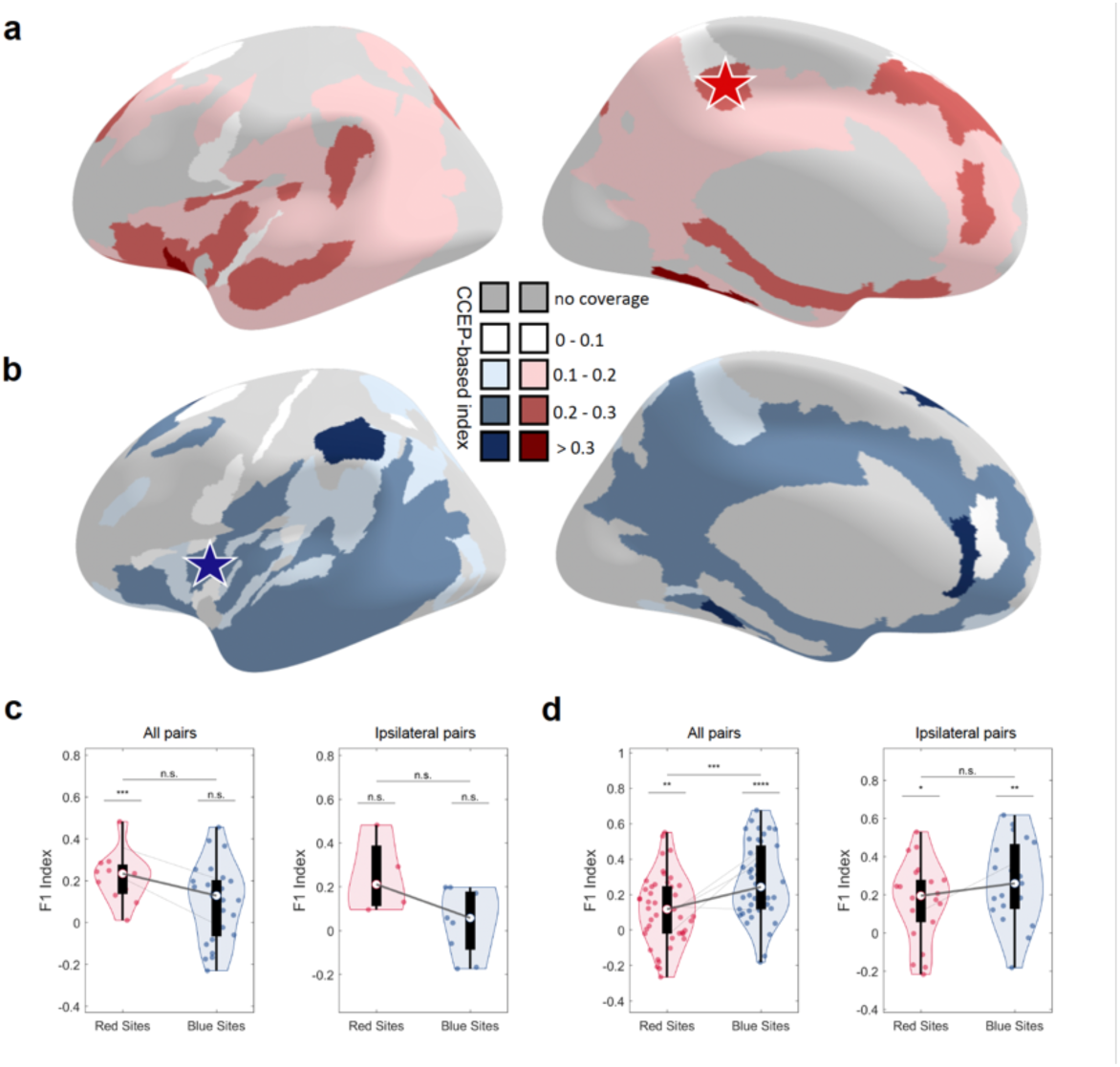
Causal Connections Between Cortical Regions and Red and Blue HPC Sites. (a-b) CCEP-based inflow connectivity index for each parcellation to red (a) and blue (b) sites. The darker the color, the higher the stronger cortical input to the HPC. Gray blocks indicate lack of coverage in those cortical parcels. In participants with *both* blue and red HPC sites *and* sufficient electrode coverage in cortical parcels, we found area 5Ci (red star in (a)) located in the dorsal posterior cingulate cortex and the area Id6 in the anterior insula (blue star in (b) to be differentially connected with red and blue HPC sites, respectively. (c) Stimulation of area 5Ci elicited significant activation in red hippocampal sites (LMM, F = 17.6556, p = 0.0013, Bonferroni-corrected for two electrode types and two regions), but not in blue sites (LMM, F = 6.2447, p = 0.0654) (left panel). When focusing only on ipsilateral pairs, a similar trend emerged, with inflow from area 5Ci being higher to red sites than to blue sites. (d) Stimulation of dorsal anterior insula produced significantly greater inflow to blue sites than to red sites (left panel; LMM, F = 11.0934, p = 0.0024, Bonferroni-corrected for two selected regions), with a similar trend observed when considering only ipsilateral pairs (right panel).

To test the hypothesis that blue and red sites receive unequal influence from cortical brain regions, we pooled data from participants with both blue and red sites identified in their HPC, along with simultaneous coverage in brain parcels. We mitigated the problem of sparse and non-homogenous electrode coverage across participants by analyzing data within the same individual brains. With these stringent inclusion criteria, we found two participants with coverage in Area 5ci in the posterior cingulate – known to be involved in executive functions^49^- and four participants with coverage in Area Id6 in the dorsal anterior insula – known to be involved in target detection^50^. Using within-participant data, we found that each of these two regions exhibited a clearly distinct preference for causal connections with red versus blue HPC sites.

The stimulation of area 5ci elicited evoked responses in the red sites in (Figure 5c, LMM, F = 32.3383, p = 0.0008, Bonferroni-corrected across two electrode types and two regions), but not in blue sites (LMM, F = 5.6484, p = 0.1032). In both of these individuals, F1 indices were consistently higher for red sites even in the contralateral HPC compared to blue sites in the ipsilateral HPC. By contrast, the dorsal anterior insula had preferred efferent causal connectivity with the blue sites (Figure 5d, LMM, F = 15.3271, p = 0.0004, Bonferroni-correction across 2 selected regions). At the individual level, 3 out of 4 participants showed higher F1 indices to blue than red sites.

## DISCUSSION

Our finding resolves some of the important inconsistencies in the human HPC and memory literature. First, we show that the hippocampal responses reported in the literature is the *summation* of *divergent patterns of activity* generated by distinct neuronal populations that are intermingled together. Our findings underscore the pitfalls of averaging signals across all recording electrodes without accounting for the significant functional heterogeneity that exists within an archicortical tissue whose anatomical architecture is the same along its anterior posterior axis. Second, we show that mesoscale field responses observed during encoding closely resemble those observed during recognition, suggesting shared underlying mechanisms across memory stages. Third, we report that the rise of HFA and ripple activity occurs only in a subset of HPC neuronal populations, but that the mere changes in the HFA power or ripple rates do not account for successful retrieval. Instead, successful memory retrieval is associated with the coordination and synchronization of the timing of changes (rises and falls) of activity across neuronal populations with idiosyncratic and unique electrophysiological responses. More specifically, we report that memory effects depend on coordinated interactions between the changes of power of more distributed LFA activity (red and blue sites) and power of HFA (only in red sites). In contrast, coordination based solely on phase–amplitude relationships between LFA and HFA does not predict memory success, suggesting that mesoscopic electrophysiological changes – at least in the human brain - extend beyond purely oscillatory mechanisms. Lastly, we provide causal findings in the human brain, using repeated electrical stimulations in cortical regions and recording across discrete HPC sites, complementing our correlative findings. Collectively, our findings demonstrate that the hippocampus has a “salt-and-pepper” functional organization at the millimeter scale and that memory is supported by mesoscale *temporally coordinated changes in the electrophysiological fields*.

Our findings are in line with a series of intracranial recording studies revealing a mosaic-like functional architecture within the cerebral cortex—including the frontal, temporal, and insular cortices—where interdigitated and spatially adjacent neuronal populations operate as distinct functional units^33,35,65–67^. In the present study, we extend this principle to the HPC, confirming the existence of mesoscale functional units within this archicortical structure. We found that red and blue sites were not confined to a specific HPC segment or subfield but rather distributed diffusely along the anterior–posterior axis of the structure and the two sites were distributed with equal prevalence. The close juxtaposition of the red and blue populations is noteworthy as its suggests that neighboring, but functionally specialized, populations of HPC neurons work in concert to integrate and process diverse types of information, each contributing uniquely to the HPC’s overall role. Our work is in line with prior animal studies documenting a possible cellular basis for the mesoscale functional heterogeneity^68–70^. Taken together, the distributed, yet heterogeneous, mosaic architecture of the HPC provides grounds for future studies testing the hypothesis that distinct neuronal components within the HPC are responsible for different computational subtasks of memory recognition, and that successful retrieval is only possible when such computational subtasks of memory recognition are coordinated together in time. Recognizing organizational complexity within the HPC opens a new avenue of mechanistic research to advance our understanding of how this brain region supports higher-order cognitive and behavioral processes such as memory retrieval. However, the present study is limited to a word recognition task. Whether similar patterns can be observed in other types of memory paradigms (e.g., navigation or picture recognition) remains an open question and will require further investigation in future studies. Additionally, the patients included in this study had seizure onset zones primarily located in the temporal lobe. Future studies are needed to determine whether and how pathological activity influences these hippocampal patterns.

An important finding in our study pertains to changes in LFA which spanned a broader band of lower frequencies (1–8 Hz), rather than a narrow band oscillatory theta rhythm. As shown in Supplementary Figure S4, we did not see clear oscillatory activity in a narrow theta band. However, we acknowledge that distinct oscillatory activities could have been merged with non-oscillatory aperiodic broadband spectral tilts and thus obscured in our analysis^12,19^. It is also possible that that the human HPC, due to its size and complexity of cortical afferents to it, does not exhibit purely oscillatory rhythms as clearly documented in rodent brains^59^.

While we acknowledge that the absence of evidence does not equate to evidence for the absence of oscillatory theta rhythms in the HPC, our data as presented here suggest that the heightened power of “theta activity” in some of the past literature may have indeed been due to heightened activity in a broader low frequency range - extending beyond the traditionally defined theta range – possibly reflecting a tilt in the local electrical fields - as the emerging evidence in the iEEG field suggests^19,39–41,60–62^.

Lastly, we emphasize that our iEEG recordings did not allow us to tease apart of the neuronal mechanisms leading to broadband changes in the LFP signal, and as a consequence a significant upward tilt in the exponent of the aperiodic activity. In our study, we were also unable to determine if the LFA recorded in red sites and blue sites were presumably generated by the same “generators” or they were anchored in discrete red vs. blue sites. These remain to be determined in future studies using more refined recording hardware.

Our electrophysiological findings provide causal evidence by showing that the afferent inputs to the HPC may be channeled preferentially to its functionally specific subpopulations (i.e., red or blue sites). For instance, repeated stimulation of area 5ci in the posterior medial region predominantly evoked responses in red sites while the repeated stimulation of area Id6 in the dorsal anterior insula evoked responses in the blue sites. Area 5ci is known to be important for integrating large scale executive processes^49^ and often activated during memory recognition processes, whereas area Id6 in the anterior dorsal insula is a component of action mode network ^63^ often associated with target detection^64^. As seen in Figure 3, we were able to document the presence of causal effective connections to the HPC from a large mantle of the cerebral cortex (where we had electrode coverage) but future large-scale studies are needed to map the preferentiality of many more cortical regions with blue vs red sites within the HPC.

## METHODS

### Participants and Data Acquisition

Multi-channel iEEG data were collected from 26 participants with refractory focal epilepsy. Participants were implanted with multiple sEEG electrodes to pinpoint the source of their seizures and guide surgical resection. The locations of electrodes were determined by clinical purpose, and the coverage in these participants included the hippocampus. The sEEG electrodes were needle like electrodes with evenly spaced contacts of 3 or 5 mm, with a diameter of 0.86 mm and a height of 2.29 mm. Intracranial sEEG data were recorded using the Nihon Kohden recording system with a sampling rate of 1000 Hz.

### Electrode Localization

Anatomical brain scans were acquired using a T1-weighted pulse sequence on a GE 3-Tesla SIGNA Magnetic Resonance Imaging (MRI) scanner before the implantation surgery, and post-surgery computerized tomography (CT) scans were obtained at our clinical facility to assist in the electrode localization process. Subsequently, individual CT scans were co-registered to the T1-weighted images and utilized for localizing the electrodes’ sites in BioImage Suite. The native-space cortical surface of each individual was reconstructed using FreeSurfer, during which the surface electrodes were registered to the individual Freesurfer space and affine-transformed to a MNI 305 space. Following this, the electrodes were overlaid on the surface and localized based on the Desikan Killiany atlas, confirmed by a trained neuroanatomist.

### Paradigm of Memory Experiment

Participants were recruited to perform the memory experiment at their bedside in the EMU. The tasks were programmed via Psychtoolbox (http://psychtoolbox.org/) in MATLAB and were run on either an Apple Macbook Pro or HP laptop. The laptop was placed about 70 cm in front of participants, while they were sitting up in their hospital bed.

The memory experiment consisted of a memory encoding test and two recognition tests. During the encoding test, participants were presented with a blank screen (dark gray background with a fixation cross). Then, common words were presented lasting for only 2000ms before the screen went back to blank while participants were instructed to press a button on a keypad to judge each item as either “abstract” (press “1”) or “concrete” (press “2”). In total 96 stimuli were shown to each participant (half words were abstract and half concrete). We measured the accuracy (i.e. whether the participant’s judgments are consistent with the real label) and reaction time (i.e. how long did it take to make the judgement) to each shown stimulus. If the participant did not respond within 5s, the program advanced to the next stimulus.

For each recognition test, the participants were presented with 96 items (48 were presented during the memory encoding test and 48 new items, no repeated words across two recognition tests) and were instructed to press a button on a keypad to judge each item as either a “old” word (press “1”) or a “new” one (press “2”). After each response, participants were also asked to rate their confidence level by pressing “1” for low confidence or “2” for high confidence. Similar to the encoding test, during the recognition tests, common words were presented lasting for only 2000ms before the screen went back to blank while participants had maximum of 5s to make their decision, otherwise the program advanced to the next stimulus.

### Analysis of Behavioral Data

Wilcoxon sum-rank tests were first measured the differences of reaction time (RT) between memory outcome (hit vs. miss vs. CR vs. FA), judgement confidence (high confidence vs. low confidence), and stimulus type (concrete vs. abstract) at participant level. We then measured the memory performance of immediate and delayed tests respectively, based on the d-prime value (d′ = z(hit rate) – z(false alarm rate)). In the subsequent analyses, we first focused on blocks with good memory performance (d’>0.5). A comparison analysis was then conducted on only blocks with poor memory performance (d’<0.5). Additional analysis including all data was conducted to measure the memory outcome between four conditions (hit vs. miss vs. CR vs. FA).

### Intracranial EEG Preprocessing

An in-house programmed preprocessing pipeline was implemented to remove noise from the iEEG signals. First, the raw signals were down-sampled to 1000 Hz. Then, electrodes located outside the brain were removed. Next, we applied notch filters with center frequency at 60, 120, and 180 Hz to remove power line noise. Subsequently, we identified channels with excessive artifacts to be excluded from subsequent analyses. Noisy and pathological channels were defined based on following criteria. (1) Raw amplitude larger than 5 times or less than one-fifth of the median raw amplitude across all channels; (2) Exhibited more than one spike per second (spikes defined as jumps between consecutive data points larger than 80 μV); (3) Pathological channels by the presence of pathological high-frequency oscillations (HFOs) using a recently developed algorithm^71^. Next, the signal from each site was re-referenced to the average of the non-noisy channels (common-average referencing). Finally, frequency decomposition of each re-referenced signal was performed using Morlet wavelet filtering at log-spaced frequencies between 1 and 256 Hz (59 total frequencies).

### Traditional Analysis of the Power in Specific Bands

The raw log-power spectrum was initially normalized by dividing the mean value during the entire task duration at each frequency. Subsequently, the data were baseline-corrected using a baseline window defined as -500ms to -200ms relative to the stimulus onset for each frequency. For each recoding site, we obtained the averaged spectrum across all trials, time-locked to both the stimulus onset and the participant’s responses, respectively. In addition, since the task was self-paced and reaction times (RTs) varied across and within participants, this analysis was also performed on spectra normalized by the percentage of RT for each trial. Specifically, we normalized the raw spectrum to a 0%–100% time window for each trial, with 500 evenly spaced time points created through linear interpolation.

To assess time-frequency power changes across specific group of recording sites, we conducted cluster-based permutation tests on the recording-site level baseline-corrected spectra. Clusters were identified by finding pixels in the time-frequency space where t-tests across recording sites revealed significant differences from baseline. The t-values within the identified clusters were summed. Subsequently, the spectra were circularly shifted 1000 times in frequency and time (while preserving their structure), and the maximum (and minimum) sum of cluster t-values was recorded for each iteration to create a null distribution. Clusters more extreme than 95% of the null distribution clusters were considered significant.

We evaluated power changes within two specific frequency bands: low-frequency activity (LFA, 1–8 Hz) and high-frequency activity (HFA, 50–130 Hz). For each trial, power in each band was computed as the average power during the first one-third of the normalized reaction time (RT) period, the last one-third of the normalized RT period, and the entire trial duration. Paired-sample t-tests were then performed to compare ripple rate changes between each task-related period and the pre-stimulus baseline.

Based on the observed HFA changes, recording sites were categorized as follows: sites showing a significant increase in HFA power during the post-onset period (300ms to 900ms relative to stimulus onset) were labeled as “Red sites”; those showing a significant decrease in HFA power during the pre-response period (650ms to 50ms before response) were labeled as “Blue sites”; sites exhibiting both effects were labeled as “Purple sites”; and the remaining sites without significant changes were labeled as “Grey sites.”

For group-level analysis, we applied linear mixed-effect models (LMMs) to assess band-power changes across specific group of sites, with participant included as a random effect. Subsequently, all p-values were Bonferroni-corrected for multiple comparisons.

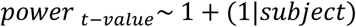

### Analysis of the Exponent

FOOOF toolbox ^41^ was utilized to calculate the aperiodic broadband exponent value at each time point. Specifically, the raw spectrum was first smoothed by a 500-ms moving window average for each frequency. Subsequently, the smoothed spectrum was interpolated into linear spaced frequencies between 1 and 256 Hz, resulting in 150 evenly distributed frequency points. The exponent at each time point was then evaluated by fitting the frequency range from 1 to 120 Hz. The exponent time series were later normalized by the mean and standard deviations (SD) of pre-stimulus baseline (-500ms to -200ms relative to the stimulus onset). For each trial, we averaged the exponent values across the entire trial duration (from stimulus onset to response) and separately calculated the average exponent during the pre-stimulus baseline (-500ms to -200ms relative to the stimulus onset). Paired sample t-tests were then performed to compare the changes of exponent from baseline for each recording site separately.

### Analysis of Sharp Wave Ripples

We measured hippocampal ripples using a method described previously^72^. Briefly, significant increases in power within the ripple band (70-180) that lasted between 20-200ms were chosen as candidate ripple events. If these events did not co-occur with inter ictal epileptic discharges (25-60Hz), they were chosen as canonical ripples.

To compare ripple rate changes during the post-onset window, pre-response, as well as the entire trial duration, we averaged the PSTH values within each respective time window and normalized by the pre-stimulus baseline (–500 ms to –200 ms relative to stimulus onset). Paired-sample t-tests were then conducted to compare ripple rate changes between each task-related period and baseline. For group-level analysis, we employed linear mixed-effects models (LMMs), including participant as a random effect. Analyses were performed across specific site groups (Red Sites, Blue Sites, Grey Sites) as well as across all hippocampal sites. All p-values were Bonferroni-corrected for multiple comparisons.

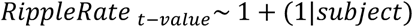

### Analysis of Pathological Activity

We first identified pathological high-frequency oscillations (pHFOs) and epileptic spikes based on previously-validated automated detection algorithms^71,73–75^. The pHFO and spike rates were then calculated for each hippocampal site by dividing the number of events by the duration of the experimental session. Finally, statistical comparisons of pathological activity across different subsets of hippocampal sites were performed using Wilcoxon rank-sum tests at the electrode level and linear mixed-effects models (LMMs) with participant included as a random effect.

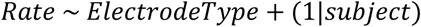

### Analysis of Response Onset Latency

We estimated the response onset latency (ROL) at each site and at the single trial level using a previously validated method ^76^. A sampling process was employed using 10,000 random windows, each of 300ms duration, to compute the average expected signal value across these windows. The averages obtained from the 300ms windows were utilized to define a permutation distribution, representing the expected average value across a 300ms window. A threshold was then established at the 95th percentile of this distribution. Significant events were identified as contiguous timepoints where the signal amplitude exceeded the threshold for more than 100ms. Notably, in red sites, we detected the positive peaks, but in blue sites we detected the negative peaks. The ROL was then defined for each of these significant events. This was computed as the average time of the event, with the calculation being weighted by the signal amplitude. For each trial, we only included the timing of the first event before subject’s response. Then, an LMM was deployed to measure the timing of ROLs between red and blues with the ROL as the dependent variable, electrode type (red vs. blue sites) as independent factors, and subject, electrode, and trials as random effects.

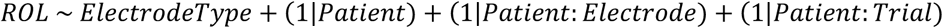

### Analysis of Phase Amplitude Coupling

We calculated phase amplitude coupling (PAC) similar to Tort et al., to examine the relationships between theta phase and HF power^77^. We extracted the phase of the theta signals ranging from 4 to 8Hz and binned the HF power (50-130Hz) in 18 uniform bins spanning -π to +π based on the post-onset time windows (300 to 900ms relative to stimulus onset) and then calculated the modulation index (MI) for each trial respectively. LMM was then performed on ipsilateral hippocampal pairs to measure the MI differences between subsets of hippocampal sites (red vs. blue).

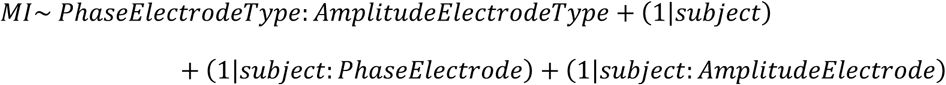

### Analysis of Functional Effects

To examine functional effects—including memory outcome (hit vs. correct rejection [CR] vs. false alarm [FA]), stimulus type (concrete vs. abstract), experimental condition (immediate vs. delayed), and judgment confidence (high vs. low)—across different subsets of hippocampal sites during recognition, we first computed key indices for each time point, including the power of specific frequency bands (e.g., LFA and HFA) and ripple rates. For each recording site, these indices were then averaged separately for specific condition time-locked to stimulus onset. To assess differences between conditions, cluster-based permutation tests with randomly shuffled labels were conducted at both the electrode levels.

To get rid of the confound effect of experimental condition (immediate vs. delayed), stimulus type (concrete vs. abstract) and judgment confidence (high confidence vs. low confidence), a linear mixed-effects model (LMM) was performed at the trial level to measure the memory outcome with experimental condition, stimulus type and judgement confidence as covariances and both participant and electrode included as random effects for each index separately. In this analysis, we only included hit and CR trials. The rational for this is that first, experimental blocks with poor performance were excluded from our analysis, resulting in an unbalanced number of trials across memory conditions—with a higher number of hit and CR trials and fewer trials in other categories. Second, the contrast between hit and CR trials has been widely used in previous studies as a reliable indicator of memory-related effects. To do this, we used the averaged indices (LFA power, HFA power, and ripple rate) within the post-onset period (300ms to 900ms relative to stimulus onset).

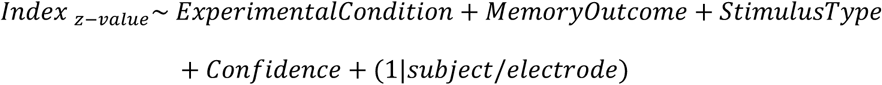

We further examined the effects of memory outcome and stimulus type by focusing exclusively on high-confidence trials, aiming to reduce potential confounding effects from familiarity, which may influence confidence judgments. Specifically, a LMM was performed at the trial level to measure the memory outcome with stimulus type as covariance and both participant and electrode included as random effects for each index separately.

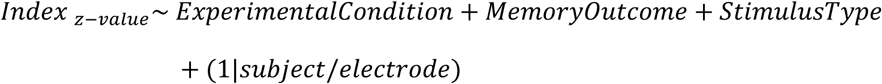

Based on the above methods, additional analyses were conducted to measure the memory outcome between four conditions (hit, miss, CR, and FA). Firstly, we identified blue and red sites using data from both immediate and delayed recognition from all 26 subjects. Subsequently, LMMs were performed at the trial level to measure the memory outcome with experimental condition, stimulus type and judgement confidence as covariances to get rid of the confound effects within red or blue sites.

To examine the correlation between changes in LFA and HFA power across red and blue sites, we analyzed nine patients who had both types of sites detected. For each trial, Pearson’s correlation was computed between the HFA power sequence at red sites and the LFA power sequence at red or blue sites from stimulus onset to response. Subsequently, a linear mixed-effects model (LMM) was used to assess differences in the correlation index (Pearson’s r) across memory outcomes (hit vs. miss vs. CR), with subject, LFA site, and HFA site included as random effects.

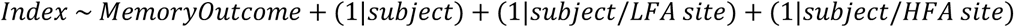

### Analysis of Cerebro-Cerebral Evoked Potentials (CCEP)

To map causal connectivity from other brain regions outside the hippocampus to the hippocampus (inflow to the hippocampus), we stimulated an adjacent pair of electrodes out of the hippocampus repeatedly (45 trials) at 6mA constant current every 2 seconds using biphasic square-wave pulses lasting 200µs while recording the iEEG signal from hippocampal sites with a sampling rate of 1000 Hz. Since the stimulations were delivered in bipolar manner across adjacent sites, we chose recording sites if either of electrodes were identified as red sites or blue sites in the hippocampus. Notably, we excluded recording site pairs if one of the pair was red, and the other was blue. In total, we obtained 5512 stimulation-recording pairs from 15 participants (2519 pairs from 11 participants were recorded in the red sites, 2993 pairs from 9 participants were recorded in the blue sites).

We used an in-house pipeline to preprocess the intracranial EEG data and calculated the connectivity index based on power spectrum and inter-trial phase coherence within the N1 period (around 20-80ms after stimulation) (F1 value in^47^). Specifically, notch filtering was performed at 60, 120, and 180Hz. Next, we excluded noisy channels. Criteria for noisy channels included extreme raw amplitude (>5 SD), frequent spikes (>3× median spike rate), and trial-level metrics such as high mean absolute voltage (>4 SD) or elevated trial variance (>3.5 SD). Trials with mean amplitudes exceeding 4 standard deviations across trials were also discarded. Then, time-frequency analysis was performed using Morlet wavelet decomposition (log-spaced from 1–256Hz, total 59 frequencies). This generated instantaneous power and phase data downsampled to 200Hz per frequency. Power spectrograms were baseline-corrected, log-transformed, and z-scored across time and frequency. Phase consistency across trials was quantified using inter-trial phase coherence (ITPC), which was square-root transformed and similarly z-scored. Manual inspection was used to validate signal integrity. We used a sliding-window cross-correlation to compare the normalized power and ITPC features with a predefined group-derived feature (F1) templates from the previous study ^47^. The F1 feature appears as a sharp wave (increased power in high gamma) with tight phase locking to the stimulation onset (strong ITPC) within the early stage after stimulation. To measure the F1 connectivity index, Pearson’s correlation coefficient was calculated across time to yield a temporal similarity curve, from which we identified the peak similarity and its timing using MATLAB’s findpeaks function. Connectivity strength was defined as the peak correlation (peak_maxCor) between stimulation–recording pairs.

To measure the strength of inflow connectivity within red sites or blue sites separately, LMMs were performed with participant and stimulation sites as random effects.

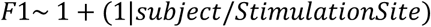

Another LMM was performed to measure the difference of inflow connectivity between red and blue sites with F1 value serving as the dependent variable and electrode type (red vs. blue) as independent factor.

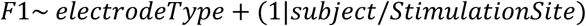

## SUPPLEMENTARY MATERIAL

### SUPPLEMENTARY RESULTS

#### Electrophysiological Analysis of Poor-Performance Blocks

We identified 12 participants who had blocks of poor performance during the delayed (d′ < 0.5) but relatively satisfactory performance (d-prime >0.5) during immediate recognition test. As detailed in the Methods and Results in the main text, we identified 34 red, 29 grey, and 26 blue sites within these cohorts using the data from the immediate memory task. Then we explored how the signature of HPC response was different in these participants during delayed memory retrieval with poor performance.

As shown in Supplementary Figure S8, red sites failed to show an increase in HFA power (LMM, F = 1.1288, p = 0.8872, Bonferroni-corrected across three site types), while blue sites failed to exhibit an increase in LFA power (F = 4.5740, p = 0.1272). Additionally, ripple rates did not significantly differ from baseline in either red (F = 2.1809, p = 0.4476) or blue sites (F = 1.1438, p = 0.8852). Additionally, we also analyzed the timing of changes in LFA and HFA across blue and red electrodes to examine if synchronization of these activities still differed as a function of memory performance in these poor-performance blocks. The correlation index between LFA in red sites and HFA in red sites (F = 0.7966, p = 0.4519), as well as between LFA in blue sites and HFA in red sites (F = 2.3006, p = 0.1003), did not differ across memory conditions within these poor-performance blocks. Similar findings were observed based on connectivity index measured by fixed time window (from stimulus onset to the median RT) (red LFA–red HFA: F = 2.23, p = 0.1081; blue LFA–red HFA: F = 0.08, p = 0.9243).

#### Additional Analysis on Memory Outcome

To further explore the memory outcome between four memory conditions (hit, miss, CR, and FA), we included data from both immediate and delayed recognition from all 26 subjects to maximize the number of samples. Using the same method as the manuscript (see Method), we identified 67 blue sites from 16 subjects and 57 red sites from 17 subjects.

As shown in Supplementary Figure S9, LMMs with experimental condition, stimulus type and judgement confidence as covariances revealed that the LFA power exhibited memory effect in both blue (F = 4.8465, p = 0.0023) and red sites (F = 4.8424, p = 0.0023). Specifically, in the blue sites, LFA power was higher in hit trials compared to CR (F = 2.914, p = 0.0107) and FA (F = 3.383, p = 0.0043). Similarly, in the red sites, LFA power was higher in the hit trials compared to CR (F = 3.570, p = 0.0022) and FA (F = 2.536, p = 0.0337).

Regarding the HFA power, the red sites did not differ between memory conditions (F = 2.4730, p = 0.0597), while the blue sites showed different HFA power across memory conditions (F = 17.896, p < 0.0001). Specifically, hit trials showed higher HFA power than CR (F = 5.976, p < 0.0001) and FA (F = 6.080, p < 0.0001). Miss trials also exhibited higher HFA power than CR (F = 2.833, p = 0.0069) and FA (F = 3.123, p = 0.0036).

Regarding the ripple rates, both blue (F = 7.5507, p < 0.0001) and red (F = 3.6381, p = 0.0122) sites exhibited memory effects. Specifically, in the blue sites, the ripple rates were significant higher in hit trials compared to miss (F = 2.339, p = 0.0463) and FA (F = 4.658, p = 0.0001). Ripple rates in CR trials also higher than those in FA trials (F = 2.590, p = 0.0384). In the red sites, hit trials showed higher ripple rates compared to CR (F = 2.403, p = 0.0463) and FA (F = 2.888, p = 0.0234).

To eliminate the confidence effect, we focused only on high-confidence trials and reconducted LMMs to measure the memory outcome. As a result, LFA power (F = 4.1337, p = 0.0062), HFA power (F = 9.0999, p < 0.0001), and ripple rates (F = 3.4699, p = 0.0154) in the blue sites still exhibited memory effects. However, LFA power (F = 2.4472, p = 0.0619) and HFA power (F = 1.6620, p = 0.1729) in the red sites did not differ between memory conditions. Ripple rates in the red sites still exhibit a weak memory effect (F = 2.7657, p = 0.0403).

In conclusion, these results further demonstrated the blue sites rather than red sites had distinct changes in activity of LFA, HFA power, and ripple rates across different memory conditions, specifically for high-confidence judgement trials.

Additionally, we further validated the *coordinated timing of activity* across LFA and HFA in red and blue sites in four memory conditions (Supplementary Figure S10). Consistent with our previous finding, the correlation coefficient index within the entire trial window between HFA in red sites and LFA in blue sites showed a significant effect of memory outcome (LMM, F = 7.3061 p = 0.0001). Specifically, the index for hit trials was significantly higher than that for miss (t = 4.025, p = 0.0002), CR (t = 2.585, p = 0.0209), and FA (t = 4.079, p = 0.0002) trials. Similarly, the index between LFA in red sites and HFA in red sites also varied across memory outcomes (LMM, F = 7.2018 p <0.0001), with hit trials (t = 3.387, p = 0.0022), miss trials (F = 4.432, p < 0.0001) and CR trials (F = 2.309, p < 0.0423) showing significantly higher indices than FA trials. By contrast, the correlation index between LFA in red sites and LFA in blue sites did not exhibit memory outcome (F = 0.5631, p = 0.6400).

## SUPPLEMENTARY FIGURES

**Figure S1:**
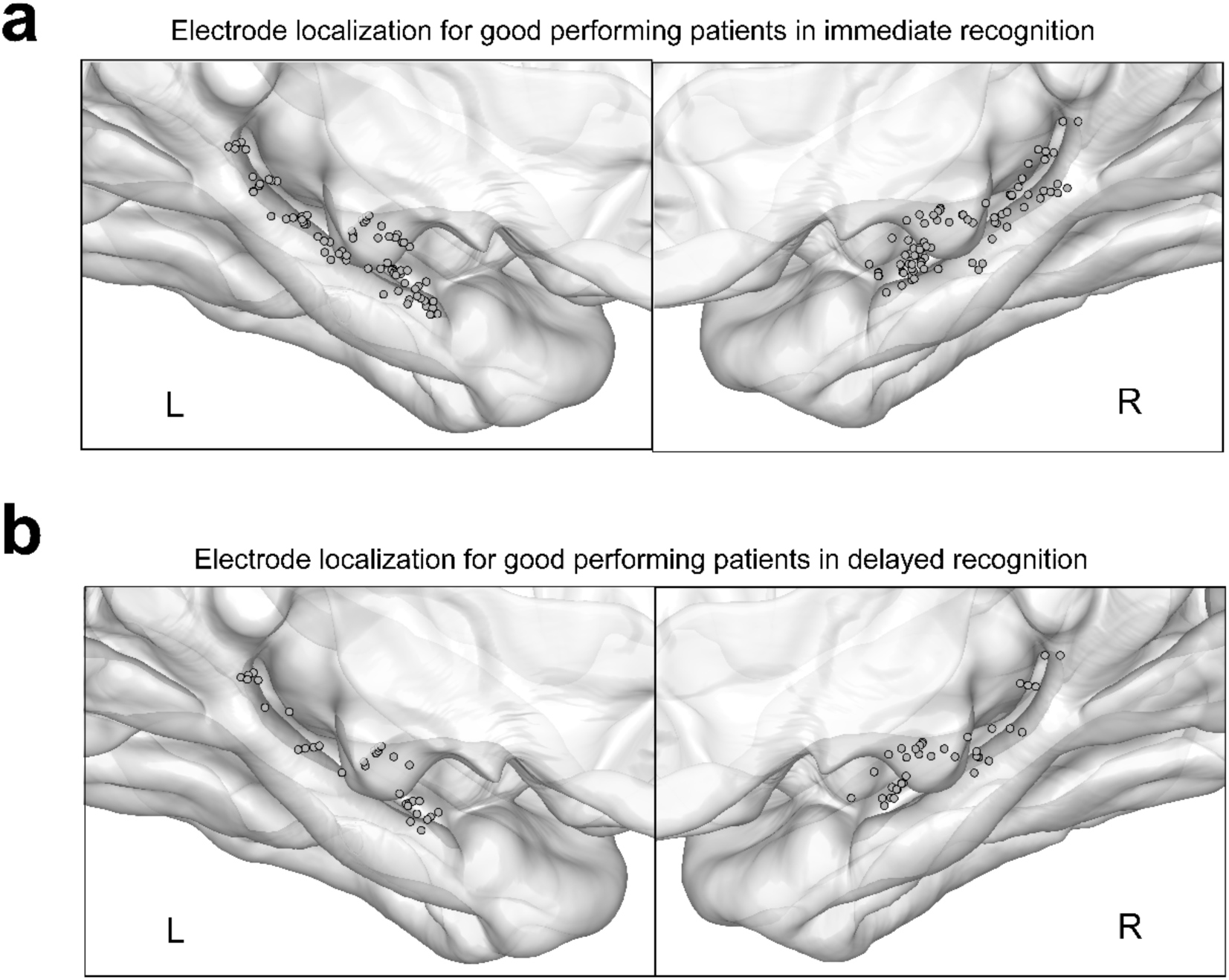
Electrode localization in the Hippocampus. (a) Electrode localization of 161 HPC sites across 22 good performing participants during immediate recognition (left hemisphere: 80 sites, right hemisphere: 81 sites). (b) Electrode localization of 71 HPC sites across 10 good performing participants during delayed recognition (left hemisphere: 34 sites, right hemisphere: 37 sites).

**Figure S2:**
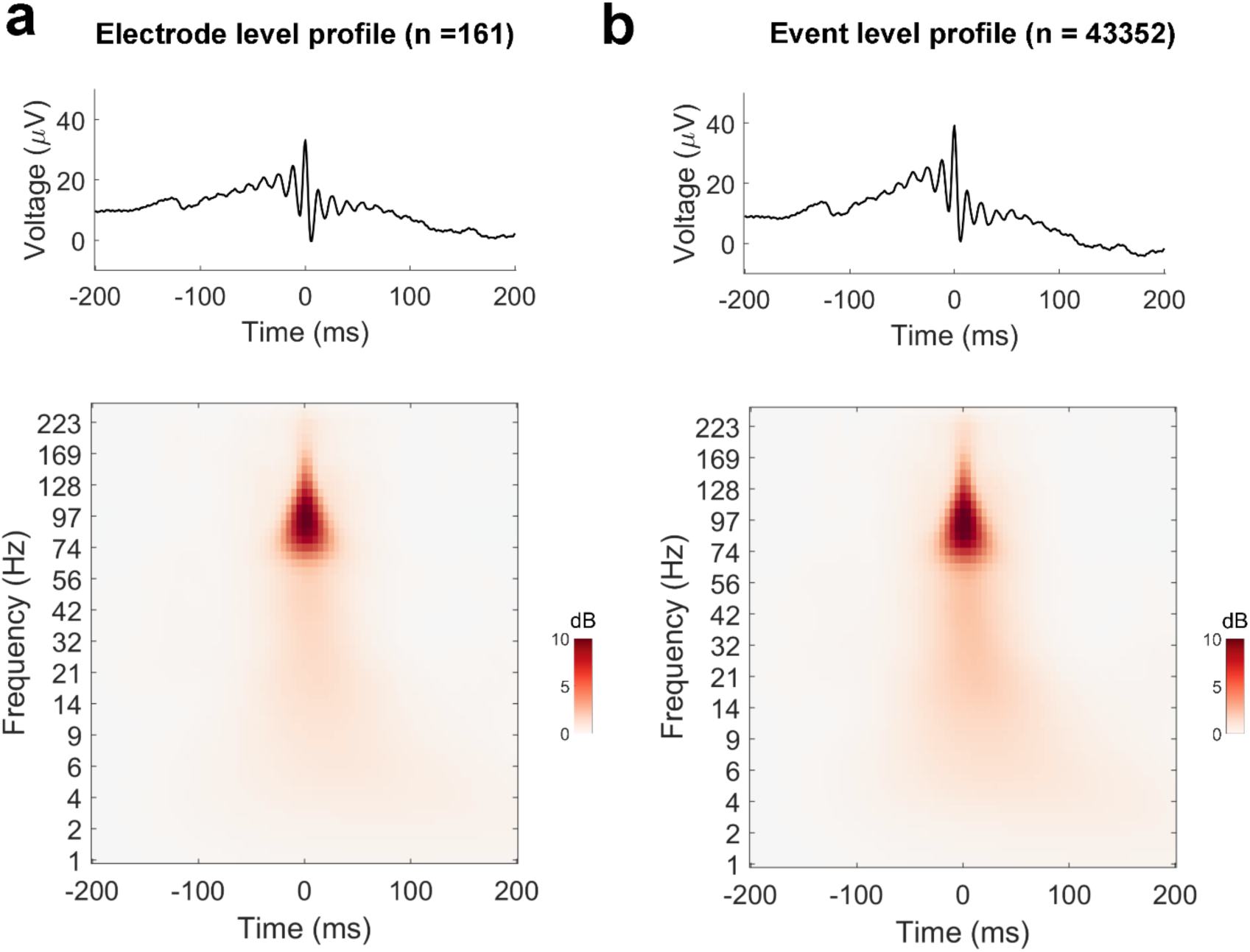
Profile of Hippocampal Sharp Wave Ripples. The averaged field potential and power spectrogram pf hippocampal ripples at electrode level (a) and across all ripple events (b).

**Figure S3:**
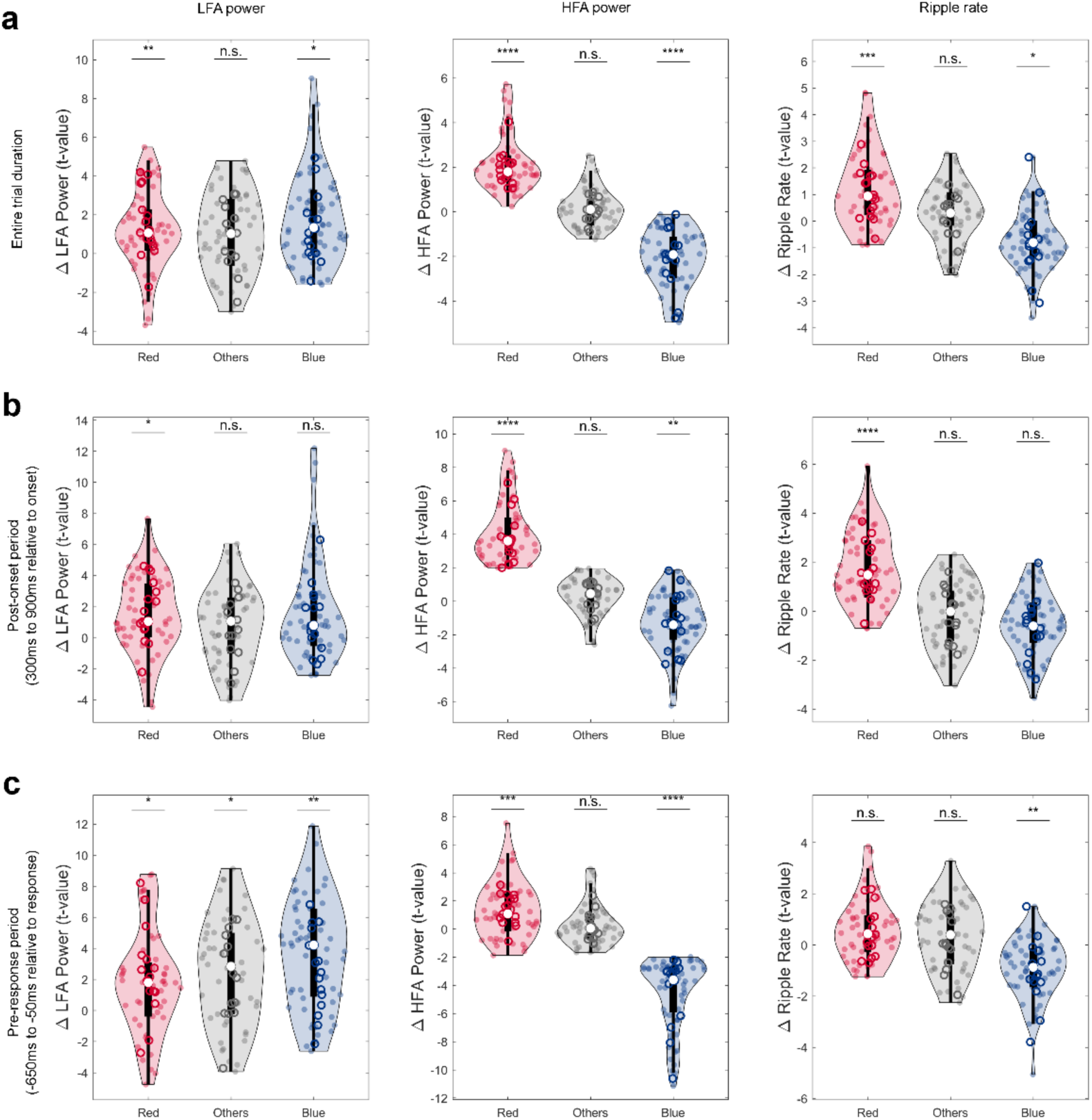
Electrophysiological Activity of Different Hippocampal Subsets. The changes of LFA power, HFA power and ripple rate during the entire trial duration (a), post-onset period (b), and pre-response period (c) from the pre-stimulus baseline (500ms to 200ms before stimulus onset) for the Red sites, Blue sites and Grey sites, respectively. **** p < 0.0001, *** p<0.001, ** p < 0.01, * p<0.05, n.s not significant, Bonferroni corrected for all p-values across three types of electrodes.

**Figure S4:**
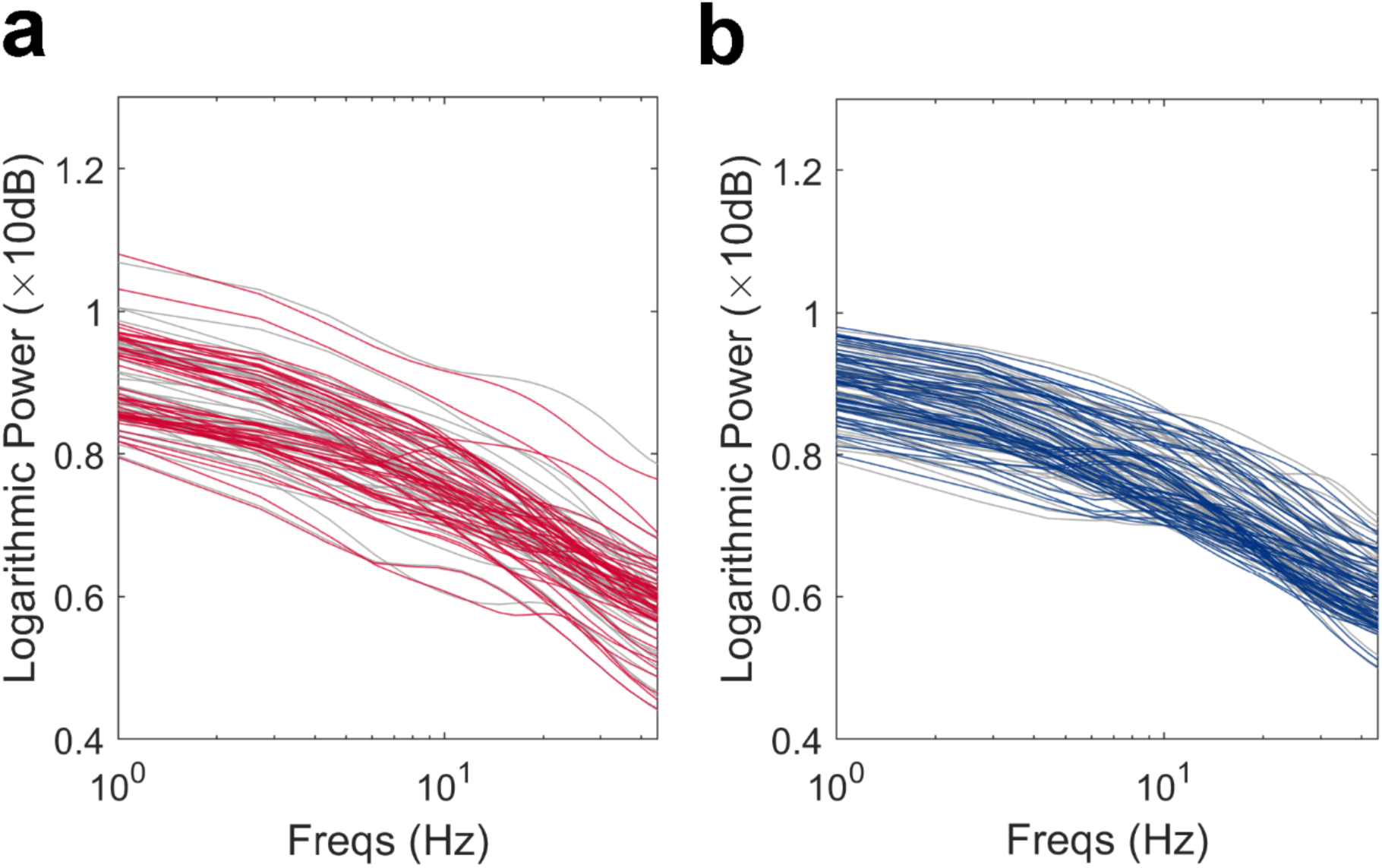
Raw Power Spectrum Within Low-Frequency Range. The raw power spectrum in log-log space did not reveal a significant oscillation with center oscillatory frequency in LFA range (1-8 Hz), which should be characterized as a “bump” above the 1/f curve, for either red sites (a) or blue sites (b) during both memory recognition (colorful lines) and pre-stimulus baseline (grey lines).

**Figure S5:**
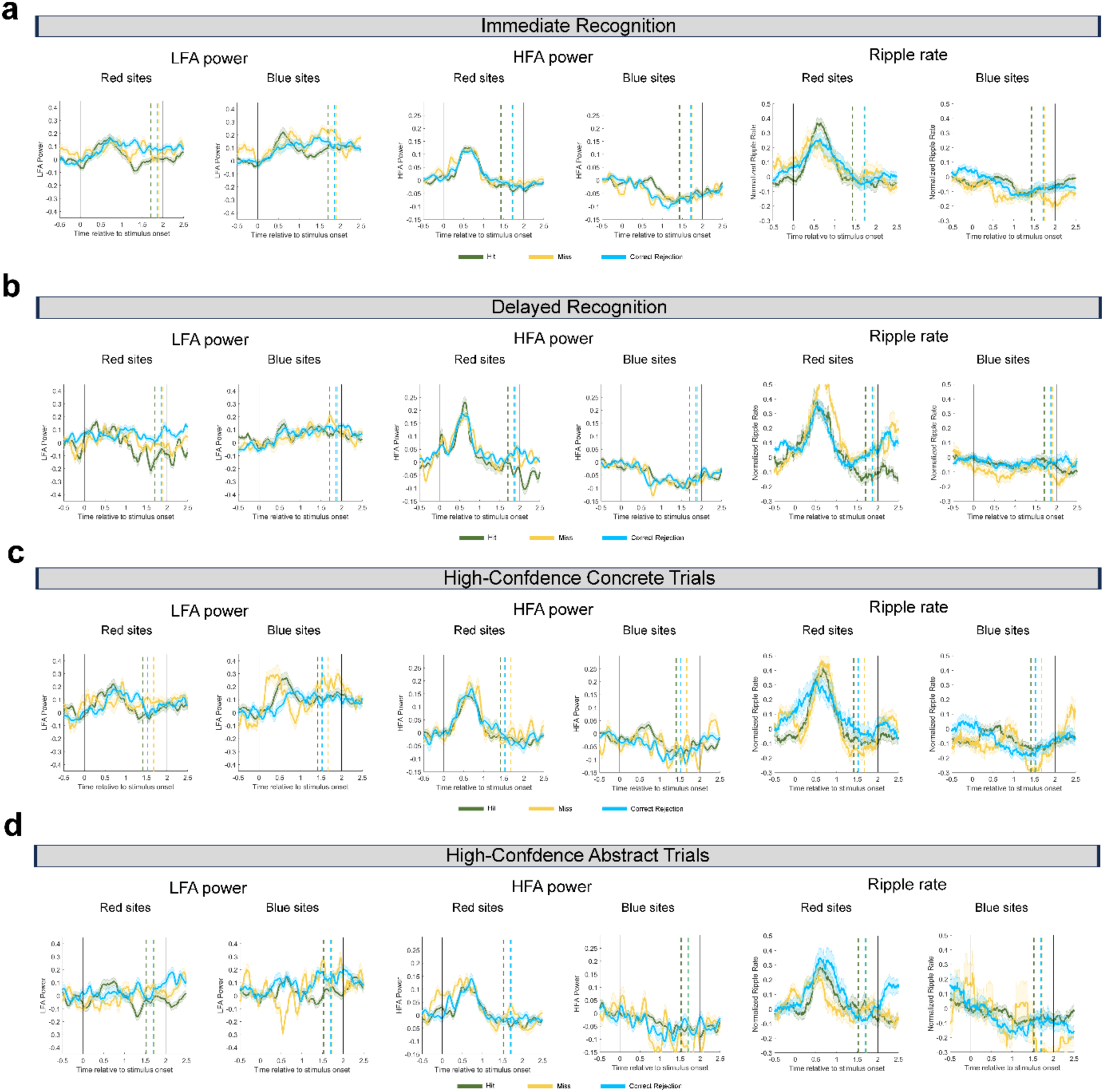
Temporal Dynamics of Signature for Memory Outcomes. Electrode-level comparisons of LFA power (left), HFA power (middle), and ripple rate (right) sequence time-locked to stimulus onset between the red sites (top) and blue (bottom) sites across different memory outcomes for (a) immediate recognition, (b) delayed recognition, (c) high-confidence concrete trials, and (d) high-confidence abstract trials.

**Figure S6:**
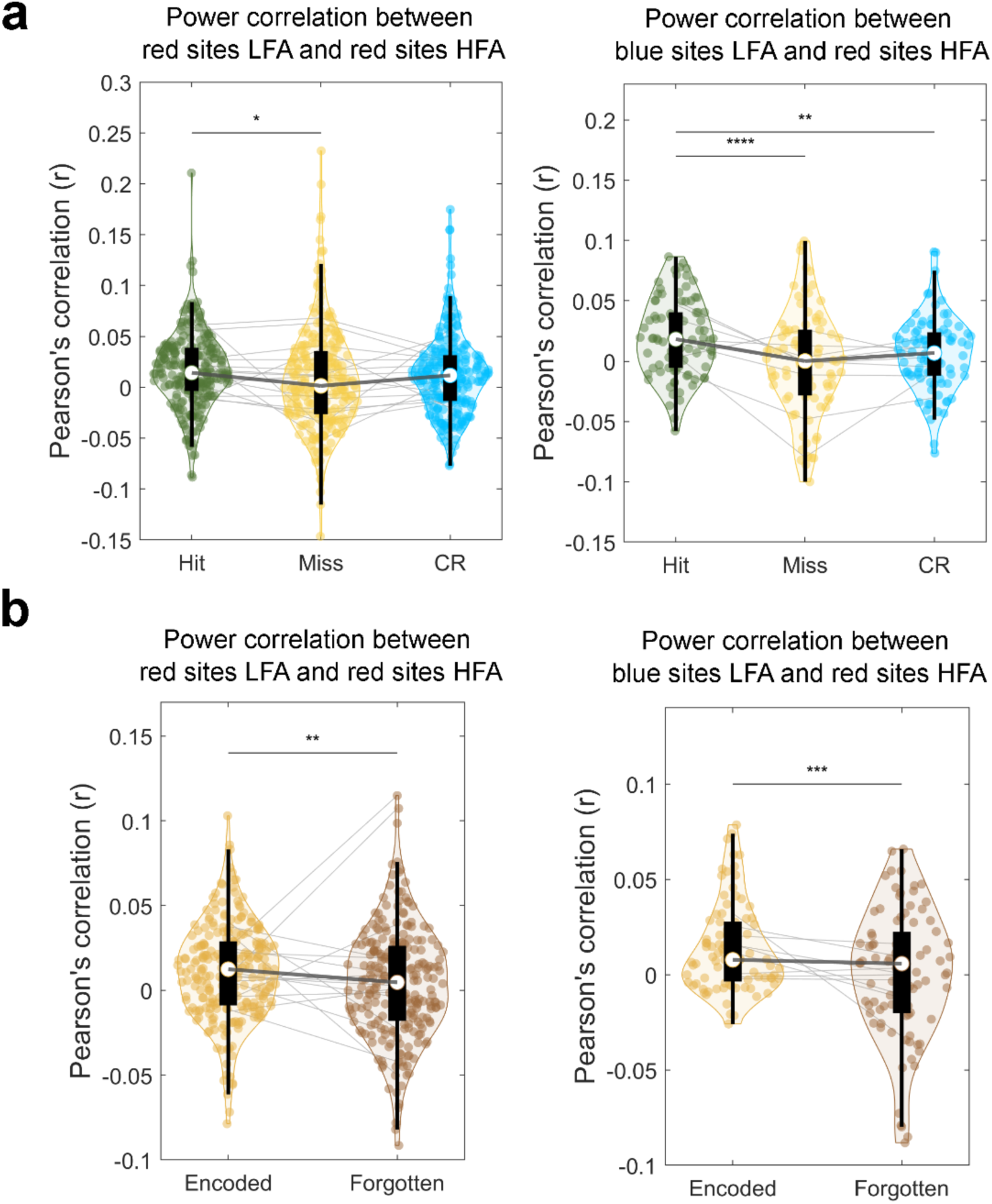
Power Correlation Between LFA and HFA. Power correlation index between LFA in (left) red sites/(right) blue sites and HFA in red sites differed between memory outcome during (**a**) recognition and (**b**) encoding.

**Figure S7:**
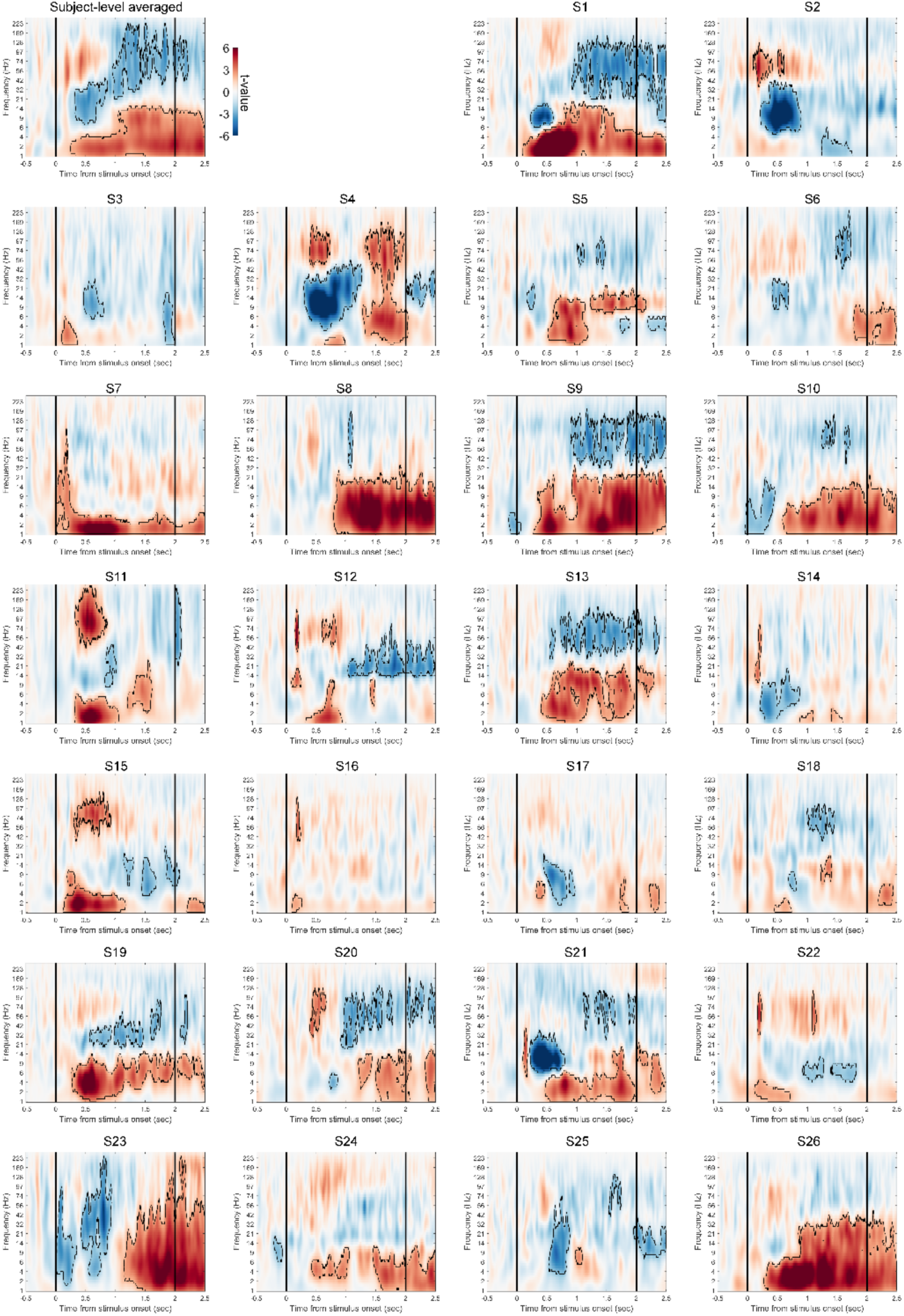
Power Spectrum During Recognition. The top-left panel shows the subject-level hippocampal (HPC) power spectrum during both immediate and delayed recognition across all 26 patients. To obtain this, power spectra were first averaged across all trials within each electrode and then averaged across all electrodes within each subject. Group-level statistics were subsequently performed at the subject level. The remaining panels illustrate trial-level power spectra across all 192 trials within each subject. For these analyses, power spectra were first averaged across all electrodes within each subject, followed by group-level statistics conducted at the trial level. In the spectrograms, time–frequency clusters showing significant power increases (red) or decreases (blue) are outlined (p < 0.05, cluster-based permutation tests). Vertical black lines indicate stimulus onset and offset.

**Figure S8:**
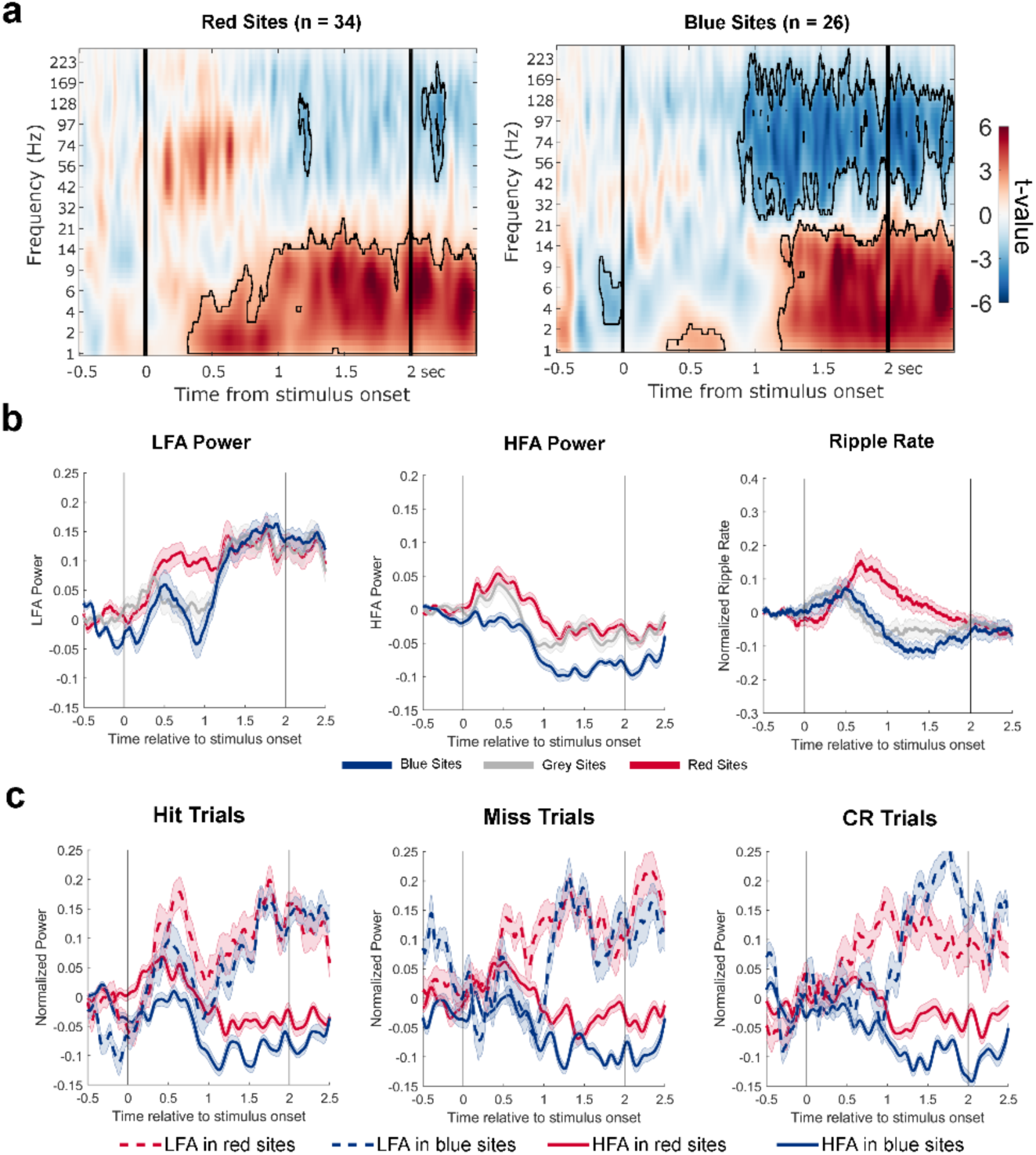
HPC Profiles During Poor-Performance Blocks. (**a**). Power spectrum of red and blue sites during poor-performance blocks from 12 subjects. In the spectrograms, time–frequency clusters showing significant power increases (red) or decreases (blue) are outlined (p < 0.05, cluster-based permutation tests). (**b**) Temporal dynamics of normalized low-frequency activity (LFA) power (left), HFA power (middle), and ripple rate (right) time-locked to stimulus onset for red, blue and grey sites. (**c**) Temporal dynamic of HFA (solid lines) and LFA (dashed lines) power across red and blue sites for hit, miss, and CR trials during poor-performance blocks. Vertical black lines indicate stimulus onset and offset.

**Figure S9:**
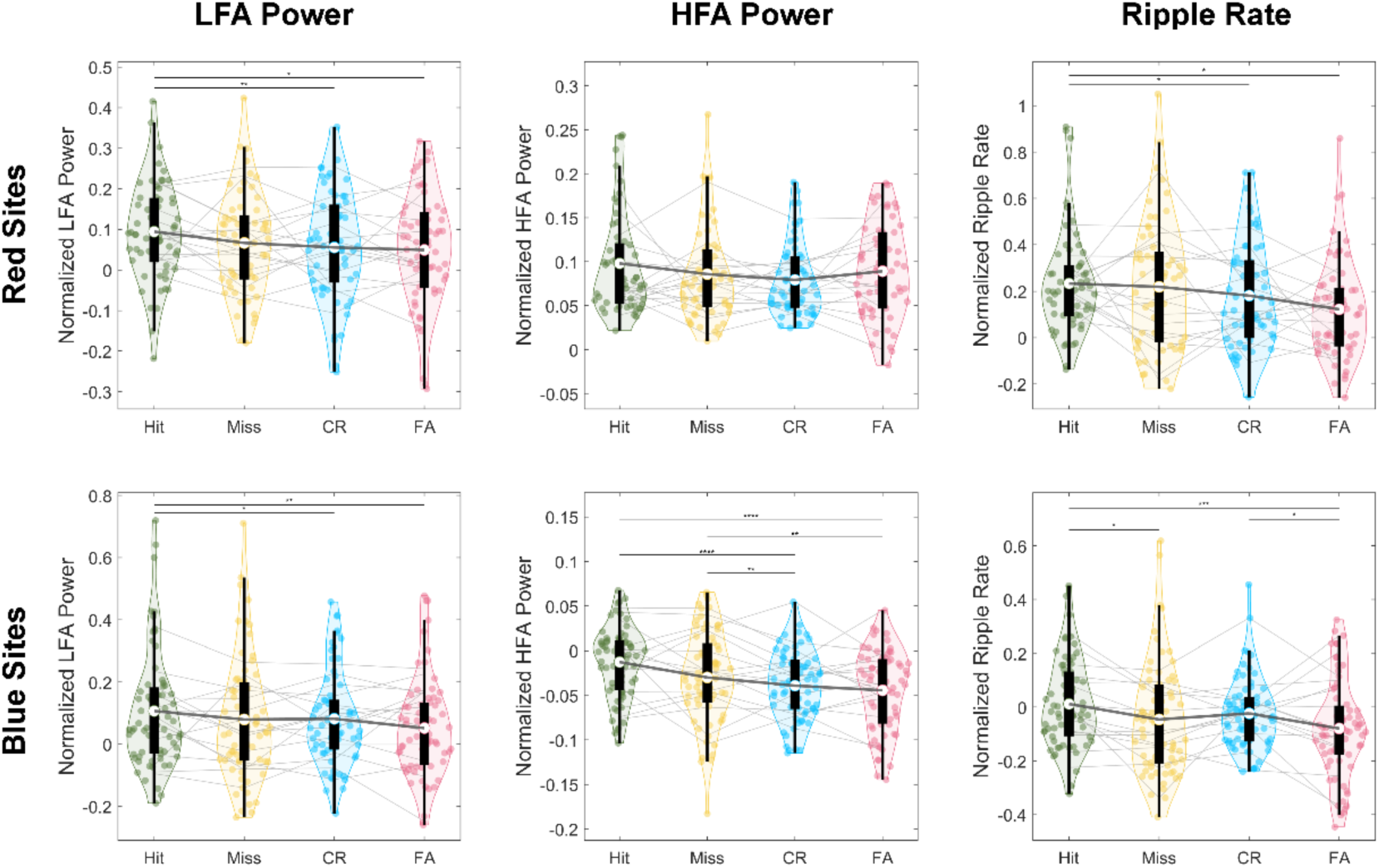
Memory Outcomes in HPC Subsets. The changes of LFA power, HFA power and ripple rate during post-onset period across memory conditions (hit, miss, CR, and FA) for the red sites and blue sites, respectively. In the violin plots, lower and upper edges of the boxplots show the 25th and 75th percentiles of the data. The white dot within the black boxes shows the mean of the data and the whiskers indicate the range within 1.5x interquartile range below 25th percentile or above 75th percentiles. The dots within the violin plot denote unique recording sites, light grey lines show individual participant averages, and dark grey lines connect the mean value across conditions. **** p < 0.0001, *** p<0.001, ** p < 0.01, * p<0.05, Bonferroni corrected for all p-values across three types of electrodes.

**Figure S10:**
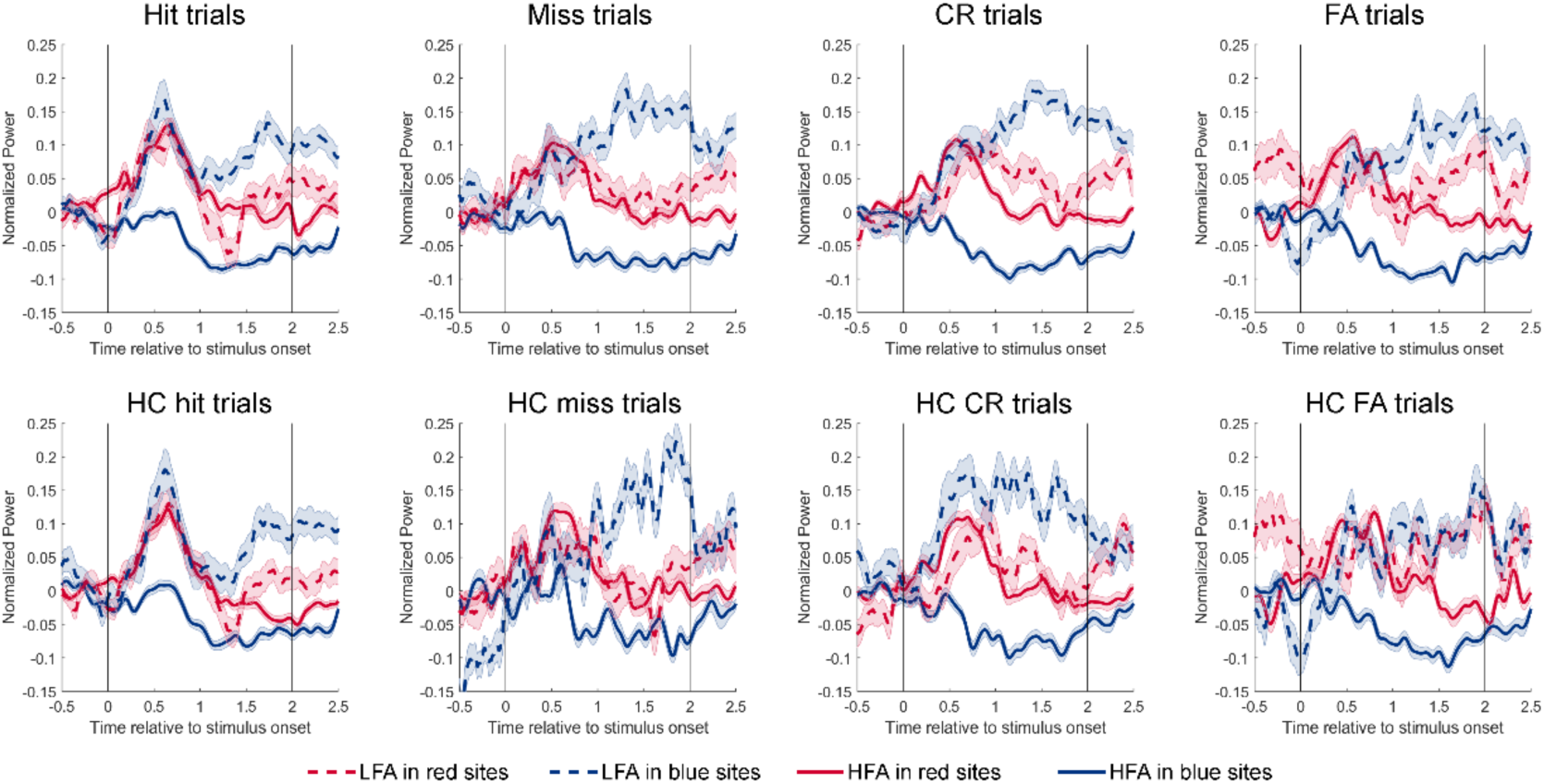
Coordination of Activity Across Four Memory Outcomes in HPC Sites. The temporal dynamic of power changes of HFA (solid lines) and LFA (dashed lines) across sites with divergent HFA responses (red and blue lines referring to red and blue sites, respectively). CR: correct rejection; FA: false alarm. HC: High confidence.

## SUPPLEMENTARY TABLES

**Table S1:** Demographics.

| Participants | Sex | Age | Race | Handedness | Language Dominance | Baseline Memory Score*<br>(0: worst;<br>15 Best Memory) | Seizures Between Recognition Tests | Medications Between Recognition Tests |
| --- | --- | --- | --- | --- | --- | --- | --- | --- |
| 1 | F | 54 | W | Right | NA | NA | No | No |
| 2 | M | 47 | H | Right | Left | NA | No | No |
| 3 | M | 35 | H | Left | Bilateral | 2.67 | Yes | No |
| 4 | F | 39 | B | Right | Left | NA | Yes | Lorazepam |
| 5 | M | 51 | A | Right | Bilateral | 9.67 | Yes | No |
| 6 | F | 37 | W | Right | NA | 11.33 | No | Midazolam |
| 7 | M | 23 | W | Right | Left | 10 | Yes | Lorazepam |
| 8 | F | 34 | W | Right | Left | 13.33 | Yes | No |
| 9 | M | 19 | W | Right | Left | 13 | Yes | Lorazepam |
| 10 | M | 28 | W | Right | Left | 11.67 | Yes | No |
| 11 | M | 33 | W | Right | Left | 12.33 | Yes | No |
| 12 | F | 36 | W | Right | Left | 7 | No | No |
| 13 | F | 35 | W | Right | NA | NA | No | No |
| 14 | M | 38 | B | Right | Left | 2.67 | No | Lorazepam |
| 15 | M | 36 | W | Right | Left | 7 | No | No |
| 16 | M | 51 | H | Right | Left | NA | Yes | Lorazepam |
| 17 | M | 57 | W | Right | Left | 13.33 | Yes | Lorazepam |
| 18 | F | 67 | W | Right | Left | 15 | No | No |
| 19 | F | 32 | ME | Right | Left | 14 | Yes | Lorazepam |
| 20 | M | 35 | H | Right | NA | 11.5 | No | No |
| 21 | F | 37 | W | Right | Left | 14.67 | No | No |
| 22 | F | 27 | B | Right | NA | NA | Yes | No |
| 23 | M | 32 | W | Right | Left | 5.67 | Yes | No |
| 24 | F | 44 | H | Right | Bilateral | 12.33 | No | No |
| 25 | F | 49 | W | Right | Left | 12.67 | No | Midazolam |
| 26 | M | 34 | W | Ambidextrous | Left | 5.67 | No | No |

\*: Measured with Ray Auditory Verbal Learning Test; A: Asian; B: Black; F: Female; H: Hispanic; M:
Male; ME: Middle Eastern; R: ethnic group based on race; W: White

**Table S2:** Performance in Memory Experiments.

| **Participant** | **Overall**<br>**D-Prime\*** | **Overall**<br>**Performance**<br>**(Immediate &**<br>**Delayed)** | **Immediate**<br>**Recognition**<br>**D-Prime** | **Immediate**<br>**Recognition**<br>**Performance** | **Delayed**<br>**Recognition**<br>**D-Prime** | **Delayed**<br>**Recognition**<br>**Performance** |
| --- | --- | --- | --- | --- | --- | --- |
| 1 | 1.43 | Good | 1.96 | Good | 1.17 | Good |
| 2 | -0.38 | Poor | -0.78 | Poor | 0.09 | Poor |
| 3 | 0.11 | Poor | 0.05 | Poor | 0.17 | Poor |
| 4 | 1.06 | Good | 1.26 | Good | 1.06 | Good |
| 5 | 1.14 | Good | 1.76 | Good | 0.64 | Good |
| 6 | 0.65 | Good | 1.16 | Good | 0.16 | Poor |
| 7 | 1.36 | Good | 2.10 | Good | 0.85 | Good |
| 8 | 0.92 | Good | 1.85 | Good | 0.32 | Poor |
| 9 | 0.67 | Good | 0.88 | Good | 0.46 | Poor |
| 10 | 1.23 | Good | 1.60 | Good | 0.90 | Good |
| 11 | 0.93 | Good | 1.46 | Good | 0.48 | Poor |
| 12 | 0.28 | Poor | 0.74 | Good | -0.17 | Poor |
| 13 | 0.84 | Good | 0.61 | Good | 1.11 | Good |
| 14 | -0.27 | Poor | 0.19 | Poor | -0.83 | Poor |
| 15 | 0.98 | Good | 1.26 | Good | 0.71 | Good |
| 16 | 0.26 | Poor | 0.61 | Good | -0.18 | Poor |
| 17 | 1.00 | Good | 1.77 | Good | 0.37 | Poor |
| 18 | 0.65 | Good | 1.18 | Good | 0.00 | Poor |
| 19 | 1.96 | Good | 2.75 | Good | 1.40 | Good |
| 20 | 0.66 | Good | 1.18 | Good | 0.21 | Poor |
| 21 | 0.73 | Good | 1.21 | Good | 0.32 | Poor |
| 22 | -0.60 | Poor | -1.06 | Poor | 0.15 | Poor |
| 23 | 0.89 | Good | 1.47 | Good | 0.55 | Good |
| 24 | 0.53 | Good | 0.64 | Good | 0.44 | Poor |
| 25 | 0.90 | Good | 1.43 | Good | 0.64 | Good |
| 26 | 0.42 | Poor | 0.54 | Good | 0.30 | Poor |

\*Measure of d-prime reflects the strength of memory: $d' = z(\text{hit rate}) - z(\text{false alarm rate})$

**Table S3:** Reaction Times In Memory Experiments.

| **Participant Index** | **Overall** | **Hit** | **Miss** | **Correct Rejection** | **False Alarm** | **High Confidence** | **Low Confidence** | **Concrete** | **Abstract** |
| --- | --- | --- | --- | --- | --- | --- | --- | --- | --- |
| 1 | 1.68 | 1.44 | 1.87 | 1.80 | 1.91 | 1.58 | 2.03 | 1.60 | 1.76 |
| 2 | 1.31 | 1.08 | 1.82 | 1.89 | 1.27 | 1.31 | 1.39 | 1.32 | 1.31 |
| 3 | 1.63 | 1.66 | 1.55 | 1.74 | 1.57 | 1.56 | 1.65 | 1.56 | 1.70 |
| 4 | 1.66 | 1.87 | 1.57 | 1.63 | 1.80 | 1.66 | 1.79 | 1.60 | 1.73 |
| 5 | 1.41 | 1.24 | 1.41 | 1.54 | 1.55 | 1.28 | 1.61 | 1.42 | 1.39 |
| 6 | 2.00 | 1.92 | 2.11 | 1.98 | 2.07 | 1.89 | 2.11 | 1.93 | 2.06 |
| 7 | 3.63 | 3.36 | 4.04 | 3.90 | 3.39 | 3.29 | 3.87 | 3.61 | 3.65 |
| 8 | 1.44 | 1.34 | 1.47 | 1.54 | 1.45 | 1.33 | 1.68 | 1.41 | 1.46 |
| 9 | 1.81 | 1.68 | 1.84 | 1.97 | 1.81 | 1.55 | 1.96 | 1.86 | 1.76 |
| 10 | 1.60 | 1.35 | 1.88 | 1.79 | 1.69 | 1.25 | 1.83 | 1.68 | 1.52 |
| 11 | 1.72 | 1.59 | 1.86 | 1.69 | 1.89 | 1.58 | 1.98 | 1.70 | 1.73 |
| 12 | 1.86 | 1.78 | 1.96 | 1.84 | 1.91 | 1.77 | 2.09 | 1.82 | 1.90 |
| 13 | 1.87 | 1.73 | 1.96 | 1.92 | 1.87 | 1.81 | 2.03 | 1.86 | 1.89 |
| 14 | 1.47 | 1.31 | 1.71 | 1.56 | 1.43 | 1.38 | 1.68 | 1.42 | 1.52 |
| 15 | 2.83 | 2.59 | 2.81 | 2.89 | 3.34 | 2.53 | 3.38 | 2.86 | 2.81 |
| 16 | 0.96 | 0.83 | 1.15 | 1.36 | 0.89 | 0.88 | 1.20 | 0.98 | 0.93 |
| 17 | 1.54 | 1.32 | 1.83 | 1.73 | 1.54 | 1.40 | 1.94 | 1.52 | 1.56 |
| 18 | 1.47 | 1.33 | 1.74 | 1.78 | 1.39 | 1.24 | 1.63 | 1.47 | 1.47 |
| 19 | 2.03 | 1.58 | 3.30 | 2.37 | 2.04 | 1.65 | 2.49 | 1.96 | 2.10 |
| 20 | 1.60 | 1.47 | 1.57 | 1.62 | 1.86 | 1.53 | 2.02 | 1.58 | 1.63 |
| 21 | 1.94 | 1.78 | 2.07 | 1.93 | 2.08 | 1.79 | 2.13 | 1.89 | 1.99 |
| 22 | 1.99 | 2.45 | 1.79 | 1.99 | 2.63 | 1.92 | 2.03 | 2.08 | 1.91 |
| 23 | 1.70 | 1.69 | 1.62 | 1.74 | 1.82 | 1.62 | 1.71 | 1.65 | 1.75 |
| 24 | 1.83 | 1.71 | 2.08 | 1.89 | 1.83 | 1.84 | 1.37 | 1.83 | 1.84 |
| 25 | 1.36 | 1.22 | 1.44 | 1.51 | 1.39 | 1.26 | 1.55 | 1.32 | 1.40 |
| 26 | 1.58 | 1.48 | 1.63 | 1.75 | 1.35 | 1.52 | 1.98 | 1.57 | 1.58 |
| Mean | 1.77 | 1.65 | 1.93 | 1.90 | 1.84 | 1.63 | 1.97 | 1.75 | 1.78 |
| SD | 0.51 | 0.51 | 0.61 | 0.50 | 0.56 | 0.46 | 0.57 | 0.51 | 0.51 |
| Median | 1.67 | 1.59 | 1.82 | 1.79 | 1.82 | 1.57 | 1.95 | 1.63 | 1.73 |

The values in the table indicate the average reaction time for each condition (second).

**Table S4:** Electrodes in Hippocampus.

| **Participant Index** | **Total Number of Hippocampal Sites** | **Red Sites** | **Grey Sites** | **Blue Sites** | **Purple Sites** |
| --- | --- | --- | --- | --- | --- |
| 1 | 13 | 2 | 0 | 8 | 3 |
| 2 | 6 | Excluded | Excluded | Excluded | Excluded |
| 3 | 6 | Excluded | Excluded | Excluded | Excluded |
| 4 | 14 | 7 | 7 | 0 | 0 |
| 5 | 3 | 1 | 0 | 2 | 0 |
| 6 | 7 | 2 | 1 | 4 | 0 |
| 7 | 6 | 0 | 4 | 2 | 0 |
| 8 | 4 | 1 | 3 | 0 | 0 |
| 9 | 18 | 3 | 5 | 10 | 0 |
| 10 | 11 | 0 | 3 | 8 | 0 |
| 11 | 8 | 4 | 4 | 0 | 0 |
| 12 | 9 | 7 | 0 | 1 | 1 |
| 13 | 2 | 0 | 0 | 2 | 0 |
| 14 | 5 | Excluded | Excluded | Excluded | Excluded |
| 15 | 4 | 3 | 0 | 0 | 1 |
| 16 | 3 | 1 | 2 | 0 | 0 |
| 17 | 6 | 1 | 5 | 0 | 0 |
| 18 | 5 | 0 | 2 | 3 | 0 |
| 19 | 7 | 2 | 4 | 1 | 0 |
| 20 | 10 | 8 | 0 | 2 | 0 |
| 21 | 7 | 0 | 2 | 5 | 0 |
| 22 | 9 | Excluded | Excluded | Excluded | Excluded |
| 23 | 6 | 0 | 3 | 3 | 0 |
| 24 | 4 | 3 | 0 | 1 | 0 |
| 25 | 5 | 1 | 0 | 4 | 0 |
| 26 | 9 | 4 | 5 | 0 | 0 |
| All | 187 | 50 | 50 | 56 | 5 |

**Table S5:** Electrode Distributions in Hippocampal Head, Body and Tail.

| Hemisphere | Hippocampal Subregion | Blue Site | Red Site | Statistics |
| --- | --- | --- | --- | --- |
| **Left** | **Head** | 14 (46.67%) | 19 (65.52%) | $\chi^2=2.1259$, $p=0.1448$ |
| | **Body** | 12 (40.00%) | 7 (24.14%) | $\chi^2=1.6993$, $p=0.1924$ |
| | **Tail** | 1 (3.33%) | 2 (6.90%) | $\chi^2=0.3879$, $p=0.5224$ |
| **Total Number** |  | 30 | 29 |  |
| **Right** | **Head** | 13 (50.00%) | 12 (57.14%) | $\chi^2=0.2381$, $p=0.6256$ |
| | **Body** | 12 (46.15%) | 1 (4.76%) | $\chi^2=9.9472$, **$p=0.0016$** |
| | **Tail** | 0 (0.00%) | 6 (28.57%) | $\chi^2=8.5157$, **$p=0.0035$** |
| **Total Number** |  | 26 | 21 |  |

The numbers in the table indicate the number of sites in each region, and the percentages in brackets indicate their proportion of the total number of sites in the corresponding hemisphere.

**Table S6:** Electrode Distributions in Hippocampal Subregions.

| Hemisphere | Subfield | Blue | Red | Statistics |
| --- | --- | --- | --- | --- |
| **Left** | **CA1** | 19 (63.33%) | 19 (65.52%) | $\chi^2=0.0307$, $p=0.8610$ |
| | **CA4/DG** | 6 (20.00%) | 6 (20.69%) | $\chi^2=0.0043$, $p=0.9475$ |
| **Total Number** |  | 30 | 29 |  |
| **Right** | **CA1** | 10 (38.46%) | 17 (80.95%) | $\chi^2=8.5800$, **$p=0.0034$** |
| | **CA4/DG** | 12 (46.15%) | 2 (9.52%) | $\chi^2=7.4529$, **$p=0.0063$** |
| **Total Number** |  | 26 | 21 |  |

The numbers in the table indicate the number of sites in each region, and the percentages in brackets indicate their proportion of the total number of sites in the corresponding hemisphere.

**Table S7:**
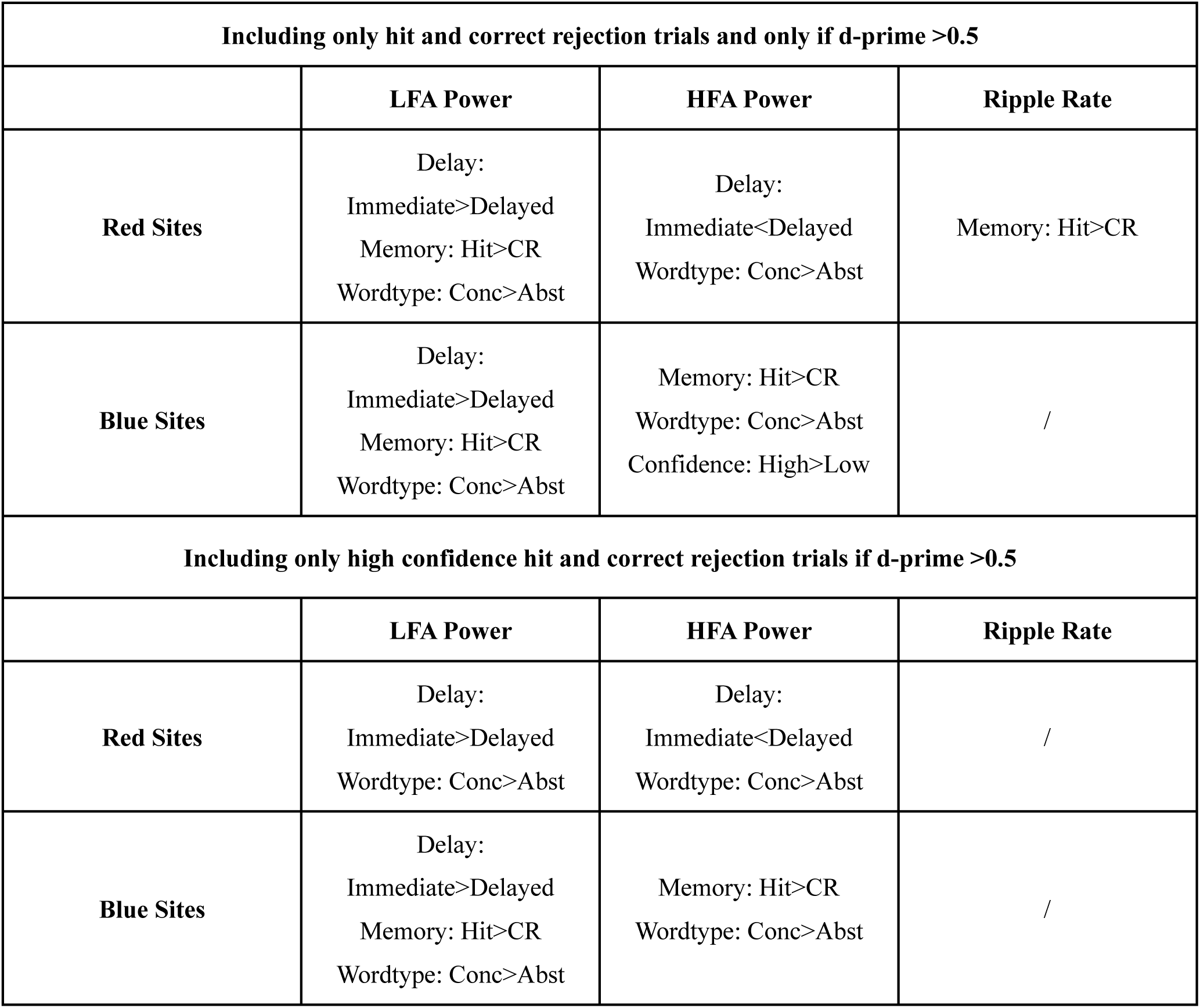
Significant Changes of Activity Across HPC Sites.

| Including only hit and correct rejection trials and only if d-prime >0.5 |  |  |  |
| --- | --- | --- | --- |
|  | LFA Power | HFA Power | Ripple Rate |
| **Red Sites** | Delay:<br>Immediate>Delayed<br>Memory: Hit>CR<br>Wordtype: Conc>Abst | Delay:<br>Immediate<Delayed<br>Wordtype: Conc>Abst | Memory: Hit>CR |
| **Blue Sites** | Delay:<br>Immediate>Delayed<br>Memory: Hit>CR<br>Wordtype: Conc>Abst | Memory: Hit>CR<br>Wordtype: Conc>Abst<br>Confidence: High>Low | / |
| Including only high confidence hit and correct rejection trials if d-prime >0.5 |  |  |  |
|  | LFA Power | HFA Power | Ripple Rate |
| **Red Sites** | Delay:<br>Immediate>Delayed<br>Wordtype: Conc>Abst | Delay:<br>Immediate<Delayed<br>Wordtype: Conc>Abst | / |
| **Blue Sites** | Delay:<br>Immediate>Delayed<br>Memory: Hit>CR<br>Wordtype: Conc>Abst | Memory: Hit>CR<br>Wordtype: Conc>Abst | / |

**Table S8:**
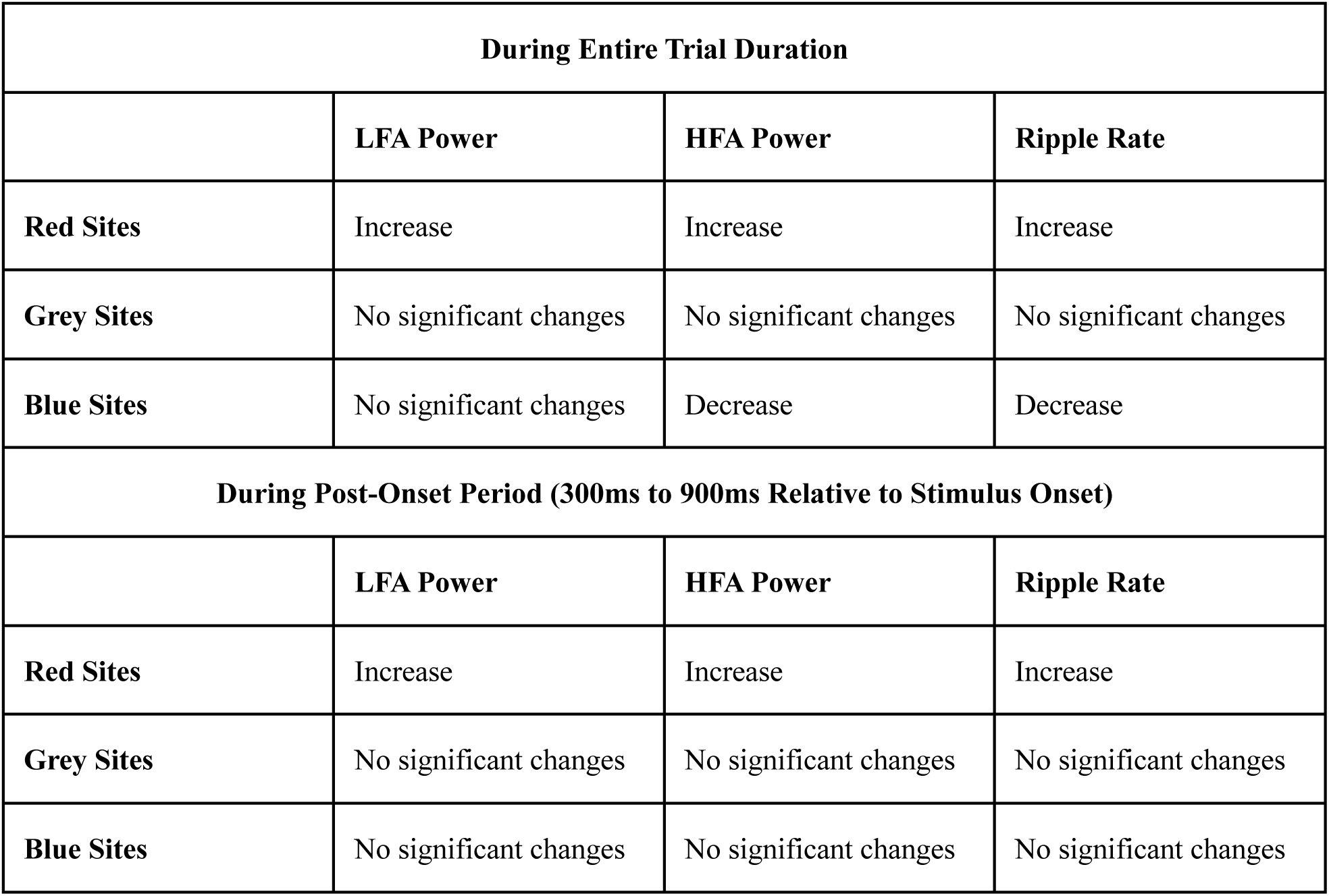
Significant Changes of Activity Compared to Pre-Stimulus Baseline for Trials with High-Confidence Judgement.

| **During Entire Trial Duration** |  |  |  |
| --- | --- | --- | --- |
|  | **LFA Power** | **HFA Power** | **Ripple Rate** |
| **Red Sites** | Increase | Increase | Increase |
| **Grey Sites** | No significant changes | No significant changes | No significant changes |
| **Blue Sites** | No significant changes | Decrease | Decrease |
| **During Post-Onset Period (300ms to 900ms Relative to Stimulus Onset)** |  |  |  |
|  | **LFA Power** | **HFA Power** | **Ripple Rate** |
| **Red Sites** | Increase | Increase | Increase |
| **Grey Sites** | No significant changes | No significant changes | No significant changes |
| **Blue Sites** | No significant changes | No significant changes | No significant changes |

## Data availability statement

Upon reasonable request, all non-identifiable data will be provided for sharing.

## Ethics approval and participant consent statement

Research procedures were approved by the IRB.

## Permission to reproduce material from other sources

NA

## Clinical trial registration

NA

